# Accessibility of Telomeric Overhangs to Stabilizing Small-Molecule Ligands

**DOI:** 10.64898/2025.12.08.692950

**Authors:** Janan Alfehaid, Vidsara Surasinghe, Sineth G. Kodikara, Mohammad L. Kabir, Masayuki Tera, Kazuo Nagasawa, John J. Portman, Hamza Balci

## Abstract

Human chromosomes terminate in 50-300 nucleotide (nt) long single-stranded telomeric overhangs composed of repeating d(TTAGGG) sequences, which can fold into tandem G-quadruplex (GQ) structures that protect chromosome ends. Stabilization of GQs by small-molecule ligands inhibits telomerase activity, motivating extensive efforts to develop GQ-targeting anti-cancer therapeutics. However, how interactions between successive GQs and bound ligands influence small-molecule accessibility remains poorly understood. Here, we employ single-molecule fluorescence microscopy and stepwise photobleaching analysis to quantify the binding stoichiometry of a fluorescently-labeled oxazole telomestatin derivative (L1Cy5-7OTD) to telomeric overhangs capable of forming 1-6 GQs (30-162 nt long), spanning much of the physiologically relevant range. We find that longer overhangs accommodate more ligands on average but exhibit consistently lower binding stoichiometry than the theoretical maximum, saturating at six molecules even in constructs with twelve binding sites. This trend was further supported by experiments showing increased L1Cy5-7OTD binding when the inter-GQ spacer was extended from 3-nt to 9-nt. This effect was independently confirmed by ensemble fluorescence enhancement experiments utilizing N-methyl mesoporphyrin IX (NMM) as a ligand. Complementary modeling with a one-dimensional lattice model describing equilibrium ligand binding to partially ordered telomeric overhangs revealed positive cooperativity between folding of successive GQs, negative cooperativity between ligands bound opposing (top and bottom) faces of successive GQs, and reduced binding affinity to GQs located at the junction of double stranded and single stranded telomeres. Together, these findings demonstrate how telomeric overhang architecture governs ligand accessibility and provide mechanistic insight to guide the rational design of GQ-targeting anticancer agents.

## Introduction

The ends of linear chromosomes resemble DNA double strand breaks (DSB) and must be protected to avoid deleterious repair and fusion. Telomeres, specialized nucleoprotein structures composed of d(TTAGGG) DNA repeat sequences and associated proteins, play an important role in this process ^1–3^. The d(TTAGGG) sequences, each called a telomeric repeat or a G-Tract, extend over 9-15 kilobases and terminate with a 50-300 nucleotide (nt) long single stranded region, called the G-tail or G-overhang ^4^. This G-overhang acts as the key loading site for telomerase, a ribonucleoprotein reverse transcriptase, which uses the free 3′-end as a primer to extend telomeres ^5–8^. Significantly, the G-overhang can fold into tandem secondary structures known as G-quadruplexes (GQs) ^9–12^. The GQ structure consists of stacked guanine tetrads, in which guanines at the tetrad corners form Hoogsteen hydrogen bonds, stabilized by monovalent cations located within or between tetrad planes ^13^. GQ formation is dynamic and can be influenced by sequence, loop length, ion concentration, and the presence of specific proteins or small-molecules ^14^. Depending on environmental conditions, the telomeric sequences can fold into parallel, antiparallel, or hybrid conformations ^15,16^. By sequestering the 3′-end of the telomere into compact structures, GQs influence telomere length regulation ^17–20^. Telomeric GQs also contribute to end protection by masking chromosome ends and preventing their misrecognition as DSB, thereby suppressing erroneous DNA damage response ^20^.

As telomerase is upregulated in most cancers and GQs can inhibit its access and catalytic activity, GQ stabilizing small-molecule ligands have emerged as potential anti-cancer therapeutics ^21–24^. Stabilization of telomeric GQs by small-molecules inhibits telomerase activity ^25–28^ and thereby reduces the proliferative capacity of cancer cells ^29,30^. Many G-quadruplex ligands feature planar aromatic scaffolds that enable strong π–π stacking onto terminal G-tetrad surfaces ^31^. Among the most potent ligands, telomestatin (from *Streptomyces anulatus*) is the first natural product shown to be a telomerase inhibitor at nanomolar concentrations by facilitating the formation or stabilization of G-quadruplex structures ^32,33^. Oxazole telomestatin derivatives (OTD) have been designed as simplified analogs of telomestatin and provide greater potential as drug candidates in terms of solubility and cytotoxicity while maintaining the GQ specificity and affinity ^34,35^. Collectively, OTDs serve as versatile chemical tools for probing GQ biology, and their strong and selective recognition of GQs also underscores their potential as scaffolds for drug development, particularly in telomerase inhibition.

To couple strong G-quadruplex recognition with direct optical detection, L1H1-7OTD molecule has been conjugated to a Cy5 fluorophore, making L1Cy5-7OTD uniquely suited for quantifying binding stoichiometry and dynamics using single-molecule fluorescence microscopy. The heptaoxazole core of 7OTD is nearly planar (dihedral angle ∼175.5°), enabling strong π–π stacking interactions with the G-quartet surface and resulting in high GQ affinity, surpassing that of related non-planar derivatives ^36^.

Using single molecule fluorescence studies, we previously reported that L1Cy5-7OTD binding to a single telomeric GQ is highly selective and stable ^37^. NMR analysis further suggested that OTDs preferentially associate with a single G-tetrad face at low OTD concentrations but are able to bind to both G-tetrads at higher concentrations ^38^. This suggests an asymmetric binding preference between the top and bottom G-tetrads of the GQ.

In addition to such fluorophore-conjugated synthetic ligands, several naturally fluorescent molecules have been employed to monitor GQ folding. Among them, N-methyl mesoporphyrin IX (NMM), displays remarkable selectivity for GQs, particularly for the parallel conformation ^39–41^. The fluorescence of NMM is strongly enhanced upon stacking onto the G-tetrads ^42^, and it preferentially binds to GQ over duplex DNA ^43^. These characteristics make NMM a sensitive probe for detecting GQ structures and interrogating their accessibility.

Despite recent advances ^44–50^, how structural features of long telomeric overhangs, such as number of G-Tracts or inter-GQ spacing, affect ligand binding and accessibility remain largely unexplored. Most prior structural studies have focused on short telomeric repeats (4-7 G-Tracts), which fold into a single intramolecular GQ. However, human telomeric overhangs are substantially longer (50–300 nt or 8-50 G-Tracts) and are capable of forming multiple (∼2-12) tandem GQs connected by flexible linkers. To address this gap, we employed single-molecule total internal reflection fluorescence (TIRF) microscopy to directly measure the binding stoichiometry of L1Cy5-7OTD on telomeric overhangs of varying lengths and structural features. In addition, we used ensemble fluorescence enhancement assay utilizing NMM to independently validate the single molecule findings. We also performed computational studies using a lattice model ^51,52^ in which the folding stability of different regions of the overhang, stacking interactions between neighboring GQs, and cooperative binding between opposing faces of successive GQs were parametrized to gain insights about the underlying structural features and interactions.

## Results and Discussion

A single molecule fluorescence assay (Fig. 1) in which the stepwise photobleaching of Cy5 fluorophores (within L1Cy5-7OTD) was utilized to determine the stoichiometry of small-molecule binding to telomeric overhangs (see Table S1 for sequences). This assay showed a concentration dependent binding stoichiometry (Fig. S1) and the optimal concentration of L1Cy5-7OTD was determined to be 1 μM. Fig. 2A shows a compilation of the photobleaching step analysis for 12 DNA constructs with varying telomeric overhang length as a 3D histogram. 2D histograms for each DNA construct are shown in Fig. S2. Only DNA constructs that showed at least one L1Cy5-7OTD binding event were included in this analysis. A similar analysis that includes the DNA constructs that did not show any binding events (zero photobleaching steps) is shown in Fig. S3. Data illustrating a potential source for the zero binding events is presented in Fig. S4. Fig. 2B and Fig. S3B show the average number of L1Cy5-7OTD molecules bound to each DNA construct. In general, the average number of bound molecules increases with overhang length, rising by ∼2.5-fold between 4GTr and 26GTr constructs. Considering that the number of GQs (1 to 6) increases by 6-fold within this length range, the observed 2.5-fold increase is modest and implies the tandem GQs do not act as independent (non-interacting) units in terms of their accessibility to L1Cy5-7OTD.

**Figure 1.**
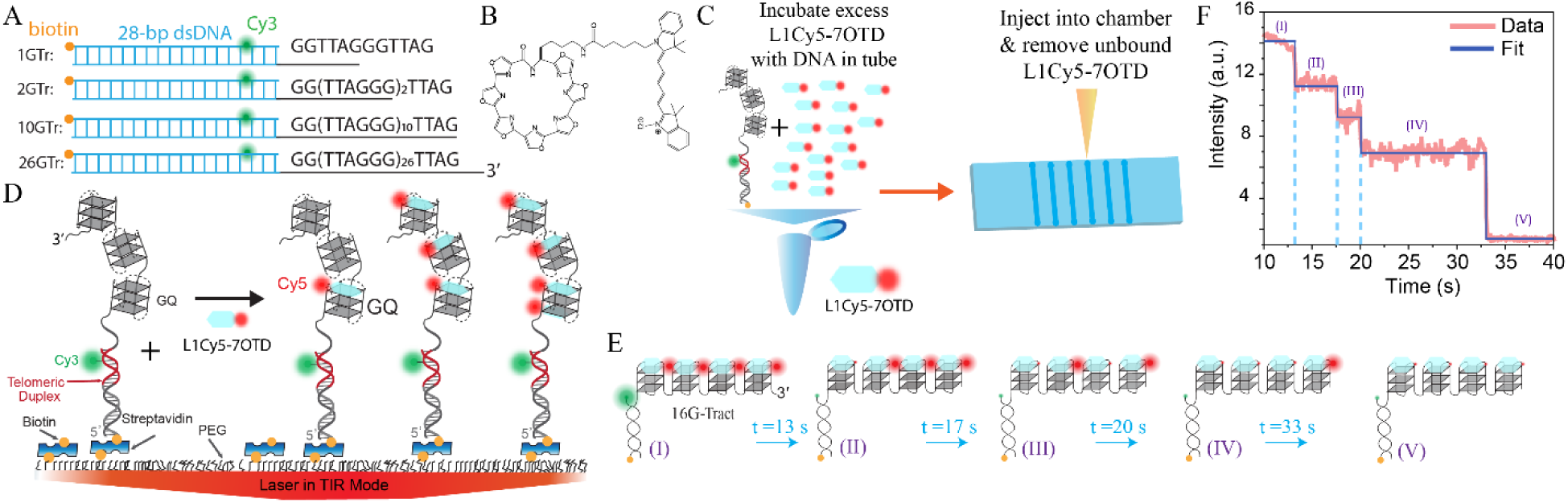
Single-molecule fluorescence studies probing L1Cy5-7OTD binding to telomeric overhangs. **(A)** The pdDNA constructs contained a 28-base pair dsDNA segment (blue) followed by single-stranded overhang of varying lengths: 1GTr, 2GTr, 10GTr, and 26GTr (where GTr is the TTAGGG repeat) are shown as examples. Cy3 was used to identify the locations of the DNA molecules with a green laser beam before it was turned off, and a red laser beam was used for stepwise photobleaching assay. Cy3 was positioned four base pairs from the duplex–overhang junction to minimize interactions with small molecule binding to GQs at that region. **(B)** Chemical structure of L1Cy5-7OTD. **(C)** L1Cy5-7OTD molecules at 1 μM concentration were mixed with partial duplex DNA (pdDNA) constructs in a tube to allow binding. After the mixture is introduced to a microfluidic channel and DNA constructs are immobilized to the surface via streptavidin-biotin linkage, the L1Cy5-7OTD molecules that are not bound to DNA were removed with a buffer exchange. **(D)** Schematic of the single-molecule TIRF microscopy assay. Cy5 molecules that co-localized with the Cy3 on the pdDNA molecules report L1Cy5-7OTD binding the GQs in the overhang. **(E)** A 16GTr construct with four tandem GQs that is initially (time interval I) bound by four L1Cy5-7OTD molecules. As time progresses (Intervals II-V), the Cy5 fluorophores sequentially photobleach, resulting in a stepwise decrease in intensity. **(F)** Fluorescence intensity trace (pink) showing the photobleaching steps depicted in (E). The data is fitted with the step finding program (blue). The four steps suggest that four L1Cy5-7OTD molecules were bound at the beginning. The labels I-V match the intervals in (E).

**Figure 2.**
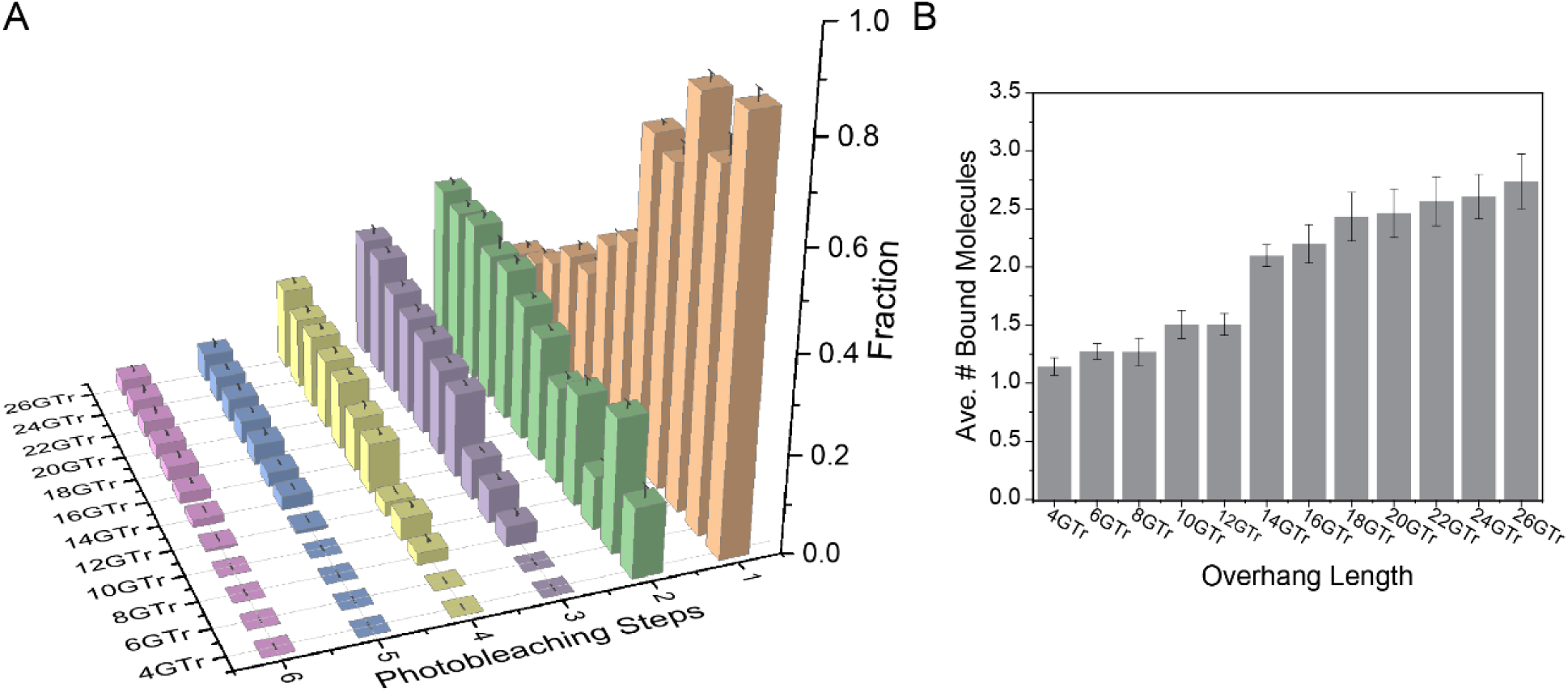
Results of photobleaching step analysis. **(A)** Three-dimensional histogram showing the normalized binding distribution of L1Cy5-7OTD molecules to 12 different DNA constructs with overhang lengths ranging from 4GTr (1 GQ) up to 26GTr (6 GQs and 2 extra G-Tracts) based on the number of discrete photobleaching steps observed in single-molecule traces. 2D versions of the histograms on each DNA construct are shown in Fig. S2. The number of DNA constructs (varied between 173-434) and photobleaching events included in each distribution (varied between 173-434) is provided in Table S2. **(B)** Average number of bound L1Cy5-7OTD molecules (〈*n*_*b*_〉) for each DNA construct. Longer overhangs exhibited higher binding stoichiometry, consistent with an increased number of binding sites. Bars represent mean values from different measurements (*n* = 5 − 10); error bars indicate the associated standard error. A one-way ANOVA revealed a significant effect of overhang length on the average number of bound molecules, *F(11, 63) = 195.42, p < .001*.

In addition to these averages, the distribution of binding stoichiometries of individual constructs can also be analyzed to deduce structural information. The 4GTr construct (one GQ) displayed a maximum of two photobleaching steps, the 8GTr construct (two GQs) showed up to four photobleaching steps, and the 12GTr construct (three GQs) reached six photobleaching steps. These results are consistent with up to two L1Cy5-7OTD molecules binding per GQ as suggested by NMR studies ^38^ which reported binding of two OTD molecules per GQ, to top and bottom G-tetrads, with significant preference to one of the tetrads. Interestingly, we did not observe more than six photobleaching steps for any of the constructs we studied, even though the fraction of molecules exhibiting six photobleaching steps continued to increase with overhang length. To illustrate, the fraction of six photobleaching steps was 6-fold higher in the 20GTr compared to the 12GTr (shortest overhang that can accommodate six L1Cy5-7OTD molecules). However, the increase in the fraction of 6-steps between 20GTr to 26GTr was within uncertainties of our measurements. We propose that this is due to stacking of adjacent GQs in longer overhangs, which induces structural compaction and thereby restricts L1Cy5-7OTD access.

Another interesting outcome of the photobleaching step analysis is the consistently higher average number of bound molecules for [4*n* + 2]GTr constructs compared to those for [4*n*]GTr, where *n* represents the maximum number of GQs the construct can form. For example, both 12GTr and 14GTr can form up to 3 GQs (*n* = 3), but the average number of bound L1Cy5-7OTD molecules (〈*n*_*b*_〉) is greater in 14GTr (2.1 ± 0.1 *vs.* 1.5 ± 0.1). This pattern is consistent across all constructs and could be due to: (i) on average more GQ molecules might form in the constructs with 4*n* + 2 G-Tracts; or (ii) the two additional repeats in [4*n* + 2]GTr constructs might function as flexible spacers and enhance ligand access to otherwise low-accessibility GQ surfaces. Interestingly, a supporting evidence for this mechanism can be deduced from a comparison of the 4GTr and 6GTr data in which only 1- or 2-step photobleaching is possible and the 6G-Tract construct demonstrates a significantly higher fraction of 2-step events compared to that of 1-step events. This particular observation cannot be explained with a greater fraction of overhangs folding into GQ, since that would have impacted the 1-step and 2-step events similarly. Instead, we propose this to be due to localization of the unfolded G-Tracts between the duplex stem and the GQ ^53^ and thereby enhancing accessibility of the GQ to small-molecules. As presented in the next section, our computational studies support this second mechanism and argue against formation of a greater number of GQs in [4*n* + 2]GTr constructs being the dominant mechanism.

To further explore this issue, we designed mutants of telomeric constructs and non-telomeric constructs in which the unfolded (spacer) regions between consecutive GQs are varied in a controlled manner. In these constructs, GQs were separated by either 3-nt d(TTA) spacers (wild-type sequence) or 9-nt d(TTATTATTA) spacers, which was implemented by replacing every fourth d(TTA) with d(TTATTATTA). Photobleaching steps analysis on these constructs revealed that constructs with 9-nt spacers consistently accommodated binding of ∼30% more L1Cy5-7OTD molecules than their 3-nt spacer counter-parts (Fig. 3A-B). Similarly, we designed non-telomeric (‘3L1L’) constructs that contained GQs formed by (GGGT)_4_ sequence, separated by 3-nt, 6-nt, or 9-nt spacers (1-3 repeats of TTA). Data on these constructs show a systematic increase of L1Cy5-7OTD binding as the spacer length (Fig. S5A) or the number of GGGT repeats (Fig. S6A) is increased. Overall, these results demonstrate that increasing the distance between GQs alleviates steric hindrance and enhances ligand accessibility.

**Figure 3.**
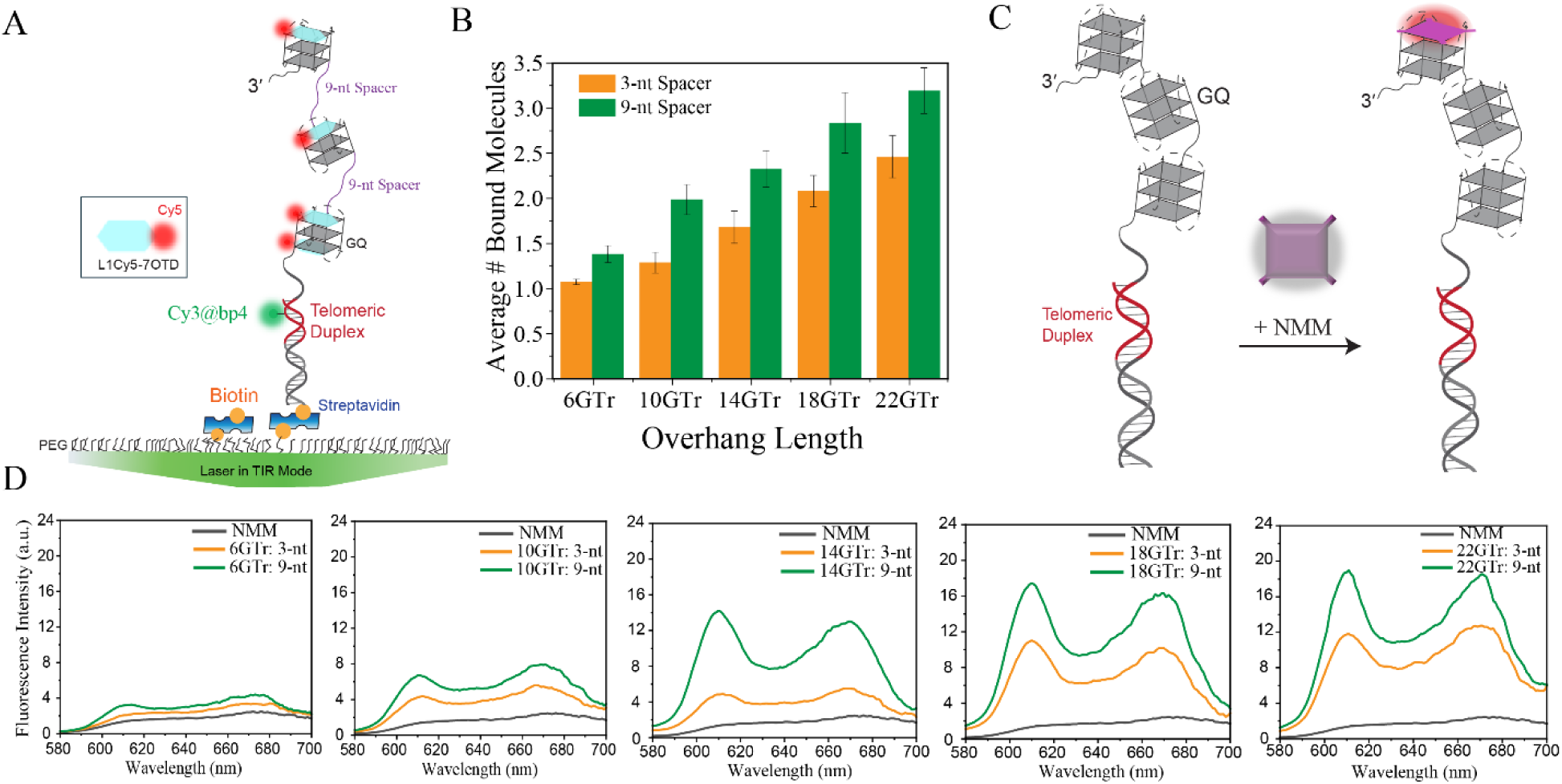
Effect of spacer length between G-quadruplexes on L1Cy5-7OTD and NMM binding. **(A)** Schematic showing DNA constructs with mutated telomeric sequences which are biased to form GQs separated by a 9 nt spacer, d(TTATTATTA), between consecutive GQs. **(B)** Average number of bound L1Cy5-7OTD molecules per DNA construct (〈*n*_*b*_〉) for overhangs with either a 3-nt or 9-nt spacer between adjacent GQs. Photobleaching step analysis was employed for quantification. Data bars represent mean values from 5–10 different measurements; error bars are the associated standard errors. The number of DNA molecules for each case (varied between 158-342) is listed in Table S3. Pairwise comparisons of each of *n*GTr construct with 3-nt or 9-nt spacer showed statistically significant differences (t-test, p<0.05 for all cases, Table S4). **(C)** Schematic of fluorescence enhancement assay in which the increase in fluorescence emission of NMM upon binding to GQ is detected. **(D)** Fluorescence spectra of NMM before (black) and after it binds to telomeric DNA constructs with 3-nt (orange) or 9-nt (green) spacers between telomeric GQs. Enhanced NMM fluorescence intensity indicates greater GQ binding and accessibility.

To independently verify GQ formation and accessibility, we employed N-methyl mesoporphyrin IX (NMM), a selective GQ probe whose fluorescence is enhanced upon binding to GQ (Fig. 3C) ^39^. Therefore, the fluorescence emission intensity of NMM can be used as a qualitative gauge for small molecule binding to telomeric overhangs. NMM emission spectra showed markedly higher fluorescence intensity in the presence of telomeric GQs compared to NMM alone (Fig. 3D). Furthermore, the NMM fluorescence emission signal was consistently greater (∼2-fold based on the peak amplitude at 610 nm) in constructs with the 9-nt spacers compared to respective constructs with 3-nt spacers (Fig. 3D). Similar experiments were performed with the non-telomeric 3L1L constructs and showed increasing NMM signal with increasing spacer length between GQs (Fig. S5B) and increasing number of GGGT repeats with a fixed 3-nt spacer (Fig. S6B).

These results demonstrate that spacer length is a critical determinant of GQ accessibility. L1Cy5-7OTD binding is determined not only by the number of GQs but also by their spatial arrangement. Constructs with 3-nt spacers between GQs likely adopt compact, stacked structures that sterically restrict access to the GQ surfaces. By contrast, longer spacers reduce these stacking interactions and expose additional binding interfaces for small-molecules. Together, these findings highlight the structural plasticity of telomeric overhangs: whereas repeat number dictates the maximum number of GQs, spacer length (unfolded regions) and stacking interactions modulate accessibility. Such structural effects are likely to be critical *in vivo*, where overhang folding heterogeneity and protein interactions may similarly control the binding efficiency of GQ ligands.

## Computational Studies

In order to better understand the implications of these experimental results in terms of underlying interactions and structural details, we performed computational studies using a transfer matrix model. This model quantifies how molecular occupancy and G-Quadruplex folding vary with telomeric overhang length. As described in Methods and Supporting Information (the Transfer Matrix section, Fig. S7 and Fig. S8), the model is parameterized by GQ folding parameters (*K*_*f*1_, *K*_f2_, *K*_*f*_, *K*_*fN*−1_ , *K*_*fN*_), nearest-neighbor cooperativity between adjacent GQs (*K*_*c*_), face-specific binding weights of small-molecules for bottom and top surfaces (*b*, *t*) with junction-proximal modifiers (*b*_1_, *t*_1_), and the cooperativity between binding to the top face of one GQ and the bottom face of the neighboring GQ (*w*_*tb*_). Fig. 4A presents a graphical summary of these parameters. The junction refers to the overhang region in the proximity of the double stranded telomere.

**Figure 4.**
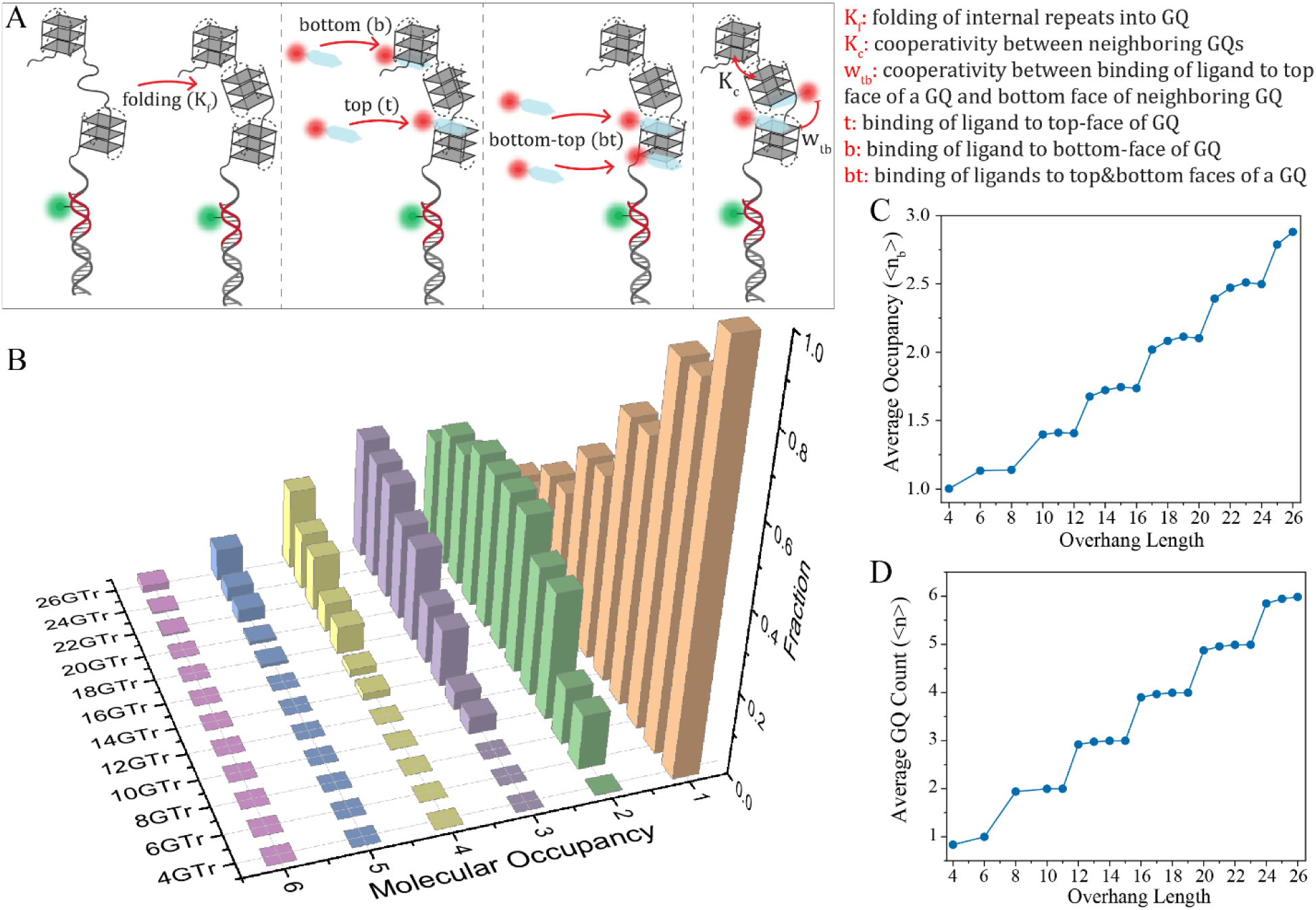
Results of the computational model calculations. **(A)** Schematics summarizing the parameters used in the model. Telomeric overhang length dependence of (**B**) probability distributions of molecular occupancy (number of bound small-molecules, *n*_*b*_); (**C**) average molecular occupancy 〈*n*_*b*_〉; and (**D**) average number of folded GQs 〈*n*〉. Normalized counts in (A) represent the likelihood of observing a given number of bound molecules across all conformational states, excluding the zero bound molecules (normalized over *n*_*b*_ ≥ 1). The distribution that includes *n*_*b*_ = 0 is presented in Fig. S9.

The calculated probability distribution of molecular occupancy, *P*_*f*_(*n*_*b*_), as a function of the number of bound molecules *n*_*b*_, represents the number of bound molecules to the overhang, is shown in Fig. 4B. This distribution is analogous to the photobleaching steps *vs.* overhang length data presented in the Fig. 2 and is similarly normalized by the weight of the states with at least one small-molecule bound (*n*_*b*_ ≥ 1). The corresponding distribution that includes overhangs with zero bound molecules is shown in Fig. S9. Our model assigns reduced junction-proximal binding parameters (*b*_1_, *t*_1_) < (*b*, *t*) to reflect lower molecular occupancy of GQs near the junction region. These assignments also reproduce the experimentally observed higher occupancy observed in [4*n* + 2]*GTr* overhangs compared to that in [4*n*]*GTr* overhangs (for a given *n*). In the special case of an overhang with only 4 G-tracts, this reduction is applied to the only GQ that can form, so the model underestimates binding compared to experiment. Although larger (*b*_1_, *t*_1_) would improve the fit for this single construct, we retain the smaller values to keep a consistent parameter set across all lengths.

For shorter overhangs, the calculated probability distributions are highly concentrated at 1-2 bound molecules; however, the number of bound molecules is significantly below the maximum possible. For example, an overhang with 12 G-Tracts can accommodate up to 6 small-molecules, yet the dominant species have 1-2 molecules. Similar to the measured distribution shown in Fig. 2, *P*_*f*_(*n*_*b*_) is a decreasing function of *n*_*b*_ for overhang lengths up to about 16GTr and develops a peak at *n*_*b*_ = 2 − 3 for longer overhangs. This molecular binding distribution constrains the values of the binding constants, *t* and *b* in the model. Incorporating the asymmetry between binding to the top and bottom interface by assuming *t* = 3 *b*, the optimum binding parameters of *b* = 0.2 and *t* = 0.6 are found. Other choices for the extent of the asymmetry are presented in Fig. S7 and Fig. S8). With higher *t* and *b* values (Fig. S10), the peak that develops in the binding distribution occurs at shorter lengths compared to the experiment (approximately at 18GTr or longer overhangs) as shown in Fig. 2 and Fig. S10. Furthermore, we found that the cooperativity between molecules bound to the top and bottom GQ surfaces of neighboring GQs, *ω*_*tb*_, is negative (i.e., *ω*_*tb*_ < 1), and that the optimal value is *ω*_*tb*_ = 0.03 for *⍺* = 3 (Fig. S7).

The binding parameters also yield an overall slope of the average molecular occupancy (〈*n*_*b*_〉), excluding zero binding events *P*_*f*_(0), as a function of overhang length that is consistent with the experimental results (Fig. 4C vs. Fig. 2B): 〈*n*_*b*_〉 systematically increases in the 1-3 range between 4-26GTr. For example, the predicted 〈*n*_*b*_〉 for 26GTr over-estimates the measured experimental average by only 4%, which is within the experimental uncertainty. Also, 〈*n*_*b*_〉 is significantly below the maximum possible *n*_*b*_, e.g., 〈*n*_*b*_〉 ≈ 3 for 26GTr which can accommodate up to *n*_*b*_ = 12 molecules. This suppression can be described with the negative cooperativity (*ω*_*tb*_ < 1) and optimal *b*, *t* values identified in the model. The calculated 〈*n*_*b*_〉 for other values of *t*, *b*, and ω_*tb*_ are shown in Fig. S11 and Fig. S12, which illustrate the typical range accommodated by the model. We note that binding cooperativity *ω*_*tb*_ is introduced in the model in an effective way to reflect steric interactions between bound molecules, without making any structural assumptions. However, loss of binding site accessibility due to three-dimensional compaction of the overhang (facilitated by positive cooperativity between successive GQs, i.e., *K*_*c*_ ≈ 5) could be the underlying mechanism for this negative cooperativity between binding to top and bottom surfaces of neighboring GQs.

The increase in 〈*n*_*b*_〉 when the overhang length extends from [4*n*]GTr to [4n + 2]GTr for a given *n* is another important consistency between the experimental observations and the model. Our model attributes this trend to unique structural and binding features of the junction region that influence both GQ folding and small-molecule binding. Interestingly, these increases occur even though the average number of GQs (〈*n*〉) remains the same for [4*n*]GTr to [4n + 2]GTr (Fig. 4D). In our previous work, similar patterns in the localization of accessible sites as a function of overhang length led us to conclude that GQ structures are significantly destabilized at the junction. That is, GQ initiated with the first or second G-Tracts exhibit reduced stability compared to those located farther from the junction (*K*_*f*1_ < *K*_*f*2_ < *K*_*f*_) ^54^. In the present work, the observed increase in 〈*n*_*b*_〉 between [4*n*]GTr and [4n + 2]GTr can be quantitatively reproduced by introducing reduced small-molecule binding constants for the GQ at the junction (*b*_1_ < *b* and *t*_1_ < *t*). To illustrate this effect, consider an overhang with [4*n*]GTr. Although there are fluctuations in folding propensity along the overhang (Fig. 5A), the most stable configuration incorporates all G-Tracts into *n* folded GQs. In contrast, a longer overhang with [4*n* + 2]GTr also tends to form *n* GQs but leaves two unstructured repeats that are more likely to localize near the junction (see Fig. 5B). Therefore, in the case of [4*n* + 2]GTr constructs, the first folded GQ is more likely to be formed by G-Tracts 3 to 6. The reduced small-molecule binding constants at the junction (*b*_1_ < *b* and *t*_1_ < *t*) decrease the 〈*n*_*b*_〉 for a [4*n*]GTr overhang but not for a [4*n* + 2]GTr overhang. In other words, the reduced *b*_1_and *t*_1_ effective leave only *n* − 1 strongly binding GQs in a [4*n*]GTr overhang. As a result, 〈*n*_*b*_〉 of [4*n*]GTr overhangs is significantly less than those of [4*n* + 2]GTr overhangs and comparable to overhangs which contain *n* − 1 GQs, i.e., [4*n* − 1]GTr, [4*n* − 2]GTr and [4*n* − 3]GTr (Fig. 4C).

**Figure 5.**
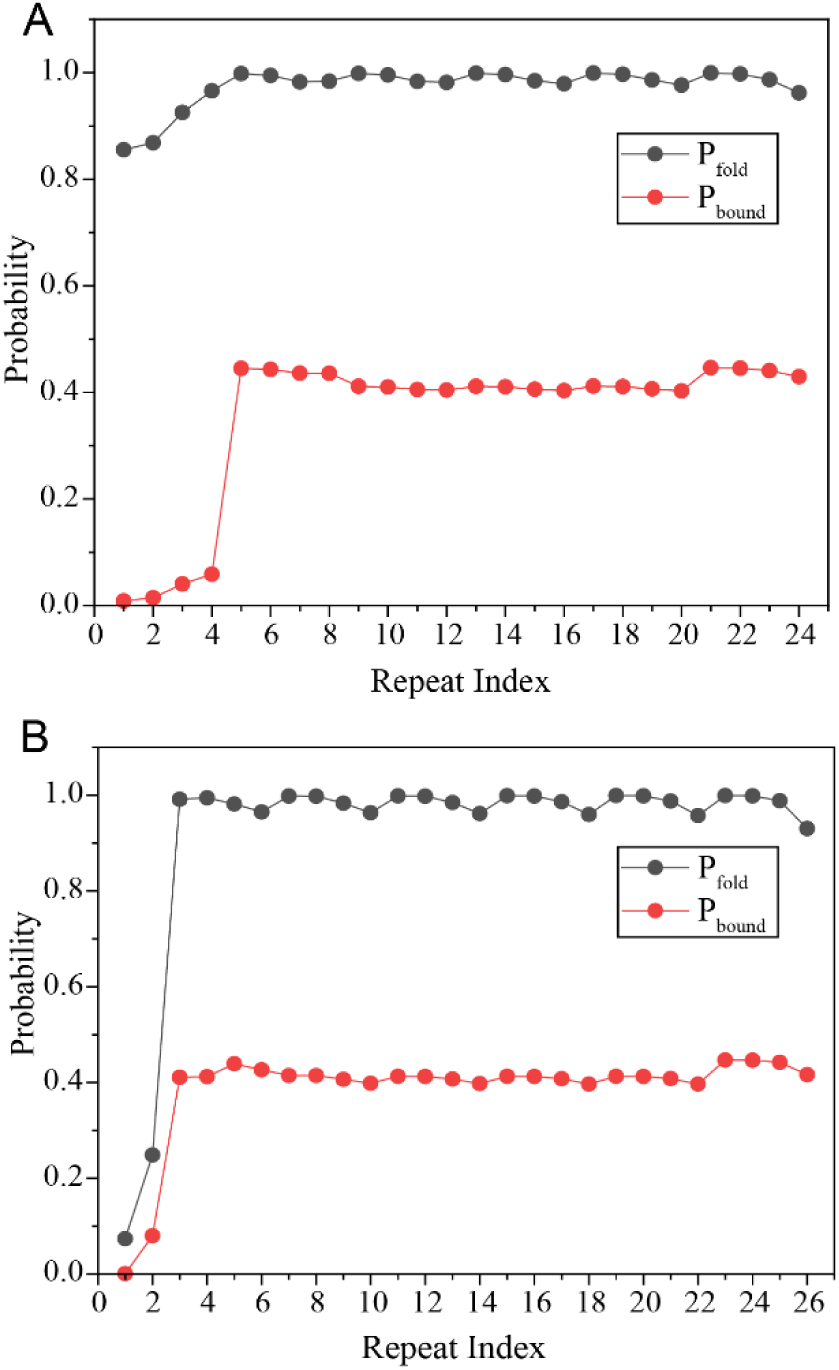
Position-dependent GQ folding and small-molecule binding profiles. The results are shown along the overhang (**A**) for 24GTr, which is an example of a [4*n*]GTr, and (**B**) for 26GTr, which an example of a [4*n* + 2]GTr. Junction destabilization, through lower GQ folding (*K*_*f*1_ < *K*_*f*2_ < *K*_*f*_), and lower small-molecule binding affinity (*b*_1_ < *b and t*_1_ < *t*) suppresses junction occupancy in [4*n*]GTr. This effectively results in *n* − 1 accessible GQs for [4*n*]GTr while all *n* GQs are accessible for [4*n* + 2]GTr. In case of [4*n* + 2]GTr, the two junction-proximal repeats are more likely to remain unfolded and separate the first GQ from the double stranded telomere, giving rise to the difference in 〈*n*_*b*_〉.

## Conclusions

This study establishes stepwise photobleaching analysis as a single-molecule approach for quantifying the stoichiometry of small-molecule binding to telomeric G-quadruplexes and introduces transfer matrix calculations as a versatile computational framework for modeling these interactions. Using L1Cy5-7OTD as a small molecule ligand, we demonstrate that binding scales with the number of GQs but remains consistently below the theoretical maximum, saturating at six molecules. Based on supporting evidence from our computational model, we attribute this behavior primarily to structural compaction and negative cooperativity between ligands bound to top and bottom faces of successive GQs. Furthermore, we show that additional G-tracts, beyond complete GQs (e.g., [4*n* + 2]GTr) act as flexible spacers, especially near the junction region, enhancing binding probability and potentially alleviating steric hindrance. Spacer-length variation supports this principle: constructs with 9-nt spacers between successive GQs consistently accommodated more L1Cy5-7OTD and NMM molecules than those with 3-nt spacers. Collectively, our findings demonstrate that telomeric overhang architecture governs ligand accessibility, providing mechanistic insight that may guide the rational design of GQ-targeting anticancer agents and telomerase inhibitors.

## Methods

### DNA Constructs

All DNA oligonucleotides (sequences are given in Table S1) were purchased from Integrated DNA Technologies (Iowa, USA) and further purified by polyacrylamide gel electrophoresis (PAGE) prior to use. Partial duplex DNA (pdDNA) constructs (Fig. 1A) were prepared by annealing a short strand (which includes a 5′-biotin and a Cy3) and a long strand which includes complementary sequences to the short strand, in addition to an overhang of repeating sequences: GG(TTAGGG)_n_TTAG, where n = 4, 6, 8, … , 26, allowing formation of 1-6 GQs. The short and long strands were mixed in 1:5 molar ratio in the presence of 150 mM KCl and 10 mM MgCl₂ for annealing. MgCl₂ concentration was later reduced to 2 mM for all experiments. A higher concentration of longer strand was used to ensure all Cy3-labeled and biotinylated short strands have a long strand partnerThe mixture was heated to 95 °C for 3 min, followed by controlled cooling to 30 °C in 1 °C steps with 3 min incubation per step using a PCR thermal cycler. Hybridized constructs for a partial duplex DNA (pdDNA) construct with a 28-bp duplex region (which includes two double stranded telomeric repeats at the junction region) and a free 3′ telomeric overhang (with varying lengths) that is available for ligand binding is obtained.

### Single Molecule Fluorescence Microscopy

All single-molecule fluorescence microscopy measurements were carried out using a prism-type total internal reflection fluorescence (TIRF) setup equipped with an Olympus IX-71 microscope and an Andor IXON EMCCD camera (IXON DV-887 EMCCD, Oxford Instruments). The IXON camera has 512 × 512 pixels and a pixel size of 16 μm. An Olympus water objective (60x, 1.20 NA) was used to collect the fluorescence signal. The total magnification in our setup is 90x, which results in an effective pixel size of 178 nm. Movies of 1000-1500 frames were acquired at 100 ms per frame (10-15 movies per construct). Imaging chambers were constructed using laser-drilled quartz slides and coverslips. The slides and coverslips were passivated with a mixture of mPEG (PEG-5000, Laysan Bio, Inc.) and biotin PEG (biotin-PEG-5000, Laysan Bio, Inc.), bonded with double-sided tape and sealed with epoxy. Before introducing the pdDNA samples, the microfluidic chamber was treated with 2% (v/v) Tween-20 to reduce non-specific binding. After 5 min incubation, the excess detergent was removed from the chamber with several successive washings and 0.01 mg/mL streptavidin was incubated for 2 minutes.

### Photobleaching Step Counting Assay

The pdDNA constructs were diluted to 400 pM in the presence of 150 mM KCl and 2 mM MgCl₂ and combined with 1 μM L1Cy5-7OTD and 10 nM quencher strand. Samples were incubated in microcentrifuge tubes for 15 min at room temperature, then introduced into microfluidic chambers and equilibrated for an additional 10 min (Fig. 1C). Excess or unbound DNA and L1Cy5-7OTD were removed by washing the channel with 20 μL of imaging buffer. The imaging buffer contains Tris-HCl (50 mM, pH 7.5), 2 mM Trolox, 0.8 mg/mL glucose, 0.1 mg/mL bovine serum albumin (BSA), 150 mM KCl, 2 mM MgCl_2_ and gloxy (glucose oxidase and catalase). In a typical smFRET experiment where highest photostability is desired, gloxy is used at 0.1-1 % concentration (1% corresponds to 0.1 mg/mL glucose oxidase, 0.02 mg/mL catalase). However, in the photobleaching step counting assay, this was reduced to 0.01% (1.0 μg/mL glucose oxidase, 0.2 μg/mL catalase) to facilitate photobleaching of fluorophores while keeping them photostable enough to resolve several consecutive photobleaching steps. The single molecule traces were analyzed using a custom MATLAB code. Statistical analysis and plots were made in Origin.

The 1 μM L1Cy5-7OTD concentration is at least an order of magnitude greater than the dissociation constant of L1Cy5-7OTD ^34,55^. Higher concentrations were avoided due to non-specific binding of L1Cy5-7OTD to the surface, which increased the fluorescence background. Measurements at lower L1Cy5-7OTD showed a concentration dependent stoichiometry (Fig. S1).

The partial duplex DNA (pdDNA) constructs are labeled with a Cy3 fluorophore, which enabled determining the locations of the DNA molecules on the surface with a green laser beam excitation (532 nm) during the first few seconds of data acquisition (Fig. 1). This was followed by turning off the green laser beam and turning on the red laser beam to image L1Cy5-7OTD molecules until all molecules photobleach (Fig. 1E-F). Only the Cy5-molecules that co-localize with a Cy3 (the surface immobilized DNA molecules) were included in analysis.

The number of L1Cy5-7OTD molecules bound to a telomeric construct was determined by counting discrete photobleaching steps, with each step corresponding to the loss of fluorescence from a single L1Cy5-7OTD molecule. Near-complete Cy5 labeling efficiency of L1Cy5-7OTD is ensured during chemical synthesis and purification. To minimize false positives from unhybridized short strands carrying Cy3, a 12-nt long quencher strand—complementary to part of the short strand and containing a broadband quencher at its 3′ end—was added. This quenching step eliminates Cy3 emission from unhybridized molecules, thereby ensuring more reliable quantification of DNA constructs that do not show L1Cy5-7OTD binding, i.e., DNA molecules with zero Cy5 photobleaching steps. Gel electrophoresis experiments suggest ∼15-30% of the short strands (with Cy3) remain unhybridized after annealing (Fig. S4). A large majority of these false-positives are eliminated by inclusion of the quencher strand ^56^, which minimally disturbs the pdDNA constructs due to already formed 28-bp long duplex segment.

### Fluorescence Enhancement Assay using NMM

N-methyl mesoporphyrin IX (NMM) was purchased from Thermo Fisher Scientific and diluted to 1 mM concentration in DMSO, followed by further dilution in an appropriate buffer. DNA constructs used for NMM studies were identical in sequence and design to those used for the L1Cy5-7OTD assay, except that the short strand lacked the Cy3 to ensure NMM is the only source of fluorescence emission. The pdDNA constructs were diluted to a concentration of 50 nM in an assay buffer containing 150 mM KCl and 2 mM MgCl₂ and mixed with 150 nM NMM in a 96-well plate with black walls and transparent bottom at room temperature. The plate was incubated in dark for 30 minutes to allow steady-state binding of NMM to the DNA constructs. Fluorescence spectra were acquired using an Agilent BioTek Synergy Neo2 plate reader with an excitation wavelength at 399 nm and emission collected from 580-700 nm. Data were analyzed by comparing the fluorescence emission of NMM alone to that of DNA-NMM complexes to quantify the fluorescence enhancement as a function of overhang or spacer length, with increased fluorescence intensity corresponding to greater NMM binding.

### Computational Model

In the absence of small molecules, the equilibrium distribution of folded GQs is described by a simple one-dimensional lattice model which describes the overhang as a sequence of *NN* TTAGGG repeats. Each repeat can either be unfolded with a statistical weight of 1 or be one of the four consecutive repeats in a folded GQ with overall statistical weight of *K*_f_. Cooperativity due to interactions between neigboring GQs is modeled with a statisical weight of *K*_c_. This model of telomeric GQ folding was introduced by Carrino and Mittermaier et al. ^57^. By fitting thermal melting data for single stranded DNA with 8 - 12 repeats (without a double stranded segment on one end), Mittermaier and coworkers reported values of *K*_f_ = 59 for interior repeats, and enhanced stabilities when GQ’s start or end (from the free ends of the ssDNA) on the last repeat, *K*_f1_ = 229, or the next to last repeat, *K*_f2_ = 80. They also reported negative cooperativity (*K*_c_ < 1) between adjacent GQs, *K*_c_ = 0.64. In our earlier work ^54^, this model was used to understand single molecule data and we found that to match localization patterns of accessible repeats, the stability for GQs at the junction region between double stranded and single stranded DNA (which were not present in Mittermaier study) needed to be significantly reduced compared to the free 3′-end, and that the cooperativity is positive (*K*_c_ > 1). Also, there was a range of parameter values consistent with the experimentally observed accessibility patterns.

While preserving the parameters that describe folding and cooperatively of GQs, this model was expanded to include small-molecule binding to GQs in the current study. As suggested by NMR studies ^38^, the model accommodated binding of two oxazole telomestatin derivative (OTD) molecules to each GQ, with a weight of *t* or *b* for binding to the top or bottom face of the GQ. Binding cooperativity of small-molecules is modeled by a statistical weight *ω*_*tb*_ associated with two molecules binding to adjacent GQs, with one molecule binding to the top interface of the downstream GQ and another binding to the bottom face of the upstream GQ.

Based on our earlier findings ^54^, we chose the following representative values to describe GQ folding for an overhang with N G-tracts: *K*_*f*_ = 60, *K*_*c*_ = 5, for the junction regions, *K*_*f*1_ = 5, *K*_*f*2_ = 10, and for the 3-prime end, *K*_*ff*_ = 229, and *K*_*ff*−1_ = 80. The free 3′-end in our constructs is equivalent to ends of the ssDNA constructs in the Mittermaier study, hence the parallel between the parameters. With the inclusion of small molecule binding, the new model fits within a class of lattice models used to describe cooperative ligand binding to one-dimensional double-stranded DNA ^58–61^. In this analogy, both folding of GQs and binding of the small molecules to the folded GQs correspond to two kinds of ligands, each with a footprint of four elementary units. Due to combinatorial frustration associated with binding at the boundaries of the DNA, patterns of binding probabilities develop with a period determined by the ligand’s footprint ^62^.

The partition function for this class of models can be calculated through transfer matrices. To do this, we denote configuration *γγ* by specifying each repeat as unfolded (1), folded (ℓ), or as folded-and-ligated. The folded and folded-ligated states are further specified based on the relative position of the repeat in the GQ and the face (top, bottom, or both) that the ligand is bound (*t*, *b* or *bt*). For example, a repeat in a GQ without a ligand is denoted by ℓ (1), ℓ (2), ℓ (3) or ℓ (4). A similar notation is used to denote a repeat in a GQ with a ligand bound to the top face (*t* (1−4)), a repeat with a bottom ligand (*b* (1−4)), and both top and bottom ligands ((*bt*) (1−4)). This notation allows us to write the partition function as:

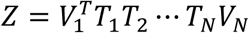

where the transfer matrix *T*_*i*_ is the conditional probability denoting the weight associated with the state of repeat *i* given the state of repeat *i* + 1. *VV*_1_ and *VV*_*f*_ are vectors enforcing allowable states of the first and last repeat. For example, the first repeat can be labeled as unfolded (1), ℓ (1), *t* (1), *b* (1), or (*bt*) (1), but it cannot be the second repeat in a GQ, ℓ (2). The matrix multiplication generates the sum of the weights of all the states in the system. The dimension of the square transfer matrix is equal to the number labels that a repeat can have in a state, 1 (unfolded) + 4 (in a GQ without a ligand) + 4 (in a GQ with only a top bound ligand) + 4 (in a GQ with only a bottom bound ligand) + 4 (in a GQ with a top and bottom bound ligands). A detailed explanation of the 17 × 17 transfer matrix for our model is given in the Supplementary Information. Many thermodynamic quantities can be directly computed with the help of transfer matrices. For example, the probability that the *i*^*th*^ repeat has particular property *β* (such as being unfolded) is

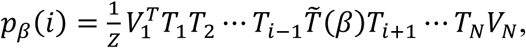

where *T̃* (*β*) is a matrix consistent with the property *β*.

Calculating the probability that there are *n*_b_ bound ligands for a construct of *NN* repeats, *P*_*f*_(*n*_b_), is not convenient with transfer matrices. Since the *P*_*f*_(*n*_b_) is proportional to the sum of the weights of configurations consistent with *n*_b_ bound ligands, we generated the weights states with transfer matrices using symbolic parameters for *t*, *b*, and *bt*, and collected appropriate terms to identify *P*_*f*_(*n*_b_) using Mathematica.

Parameters for small molecule binding are obtained by the least-squared fit method to the measured *P*_*f*_(*n*_b_) data set (see Supporting Information for details). Because experimental observations have shown that the binding affinity of OTDs to the bottom surface of the GQ is lower than that to the top surface ^38^, we investigated the influence of this asymmetry on the fit. Enforcing the binding asymmetry through the constraint *t* = *⍺b*, we find the best fit for equal binding coefficients, t = 0.4 and b = 0.4. However, fits for *⍺* > 1, are also of comparable quality (see Fig. S8). For illustration, we choose the optimal binding constants for *⍺* = 3, i.e., *t* = 0.6 and *b* = 0.2.

The cooperativity parameter ω_*tb*_ controls the distribution of ligands for larger number of repeats. We find that the optimal value of ω_*tb*_ always indicates negative cooperativity (ω_*tb*_ < 1), and for α = 3, its value is *ω*_*tb*_ = 0.03 (see Fig. S7). These parameters provide a good qualitative agreement with the measured molecular occupancy distributions. Further, to account for the special features of the junction-proximal repeats ^53,54^, binding constant *b*_1_ and *t*_1_ were introduced, where *b*_1_ < *b* and *t*_1_ < *t*. Setting *b*_1_ = 0.002 and *t*_1_ = 0.01 in the model results in a good fit with the experimental observation of an increase in small molecule binding when the number of repeats is increased from 4*n* to 4*n* + 2.

### Supporting Information Available

Sequences of DNA constructs, tables listing number of molecules included in each dataset and statistical analysis demonstrating significance of observed variations, gel image showing fraction of unhybridized short strands, experiments illustrating concentration dependence of binding stoichiometry, analysis including zero binding steps, NMM binding experiments for 3L1L constructs, description of transfer matrix model for our system, least squares fit procedure illustrating how various model parameters are obtained and the sensitivity of binding distributions to variations in these model parameters including varying *t*, *b*, and *w*_*tb*_.

## Acknowledgements

This study was supported by the National Institutes of Health under Grant No. 1R35GM156183 to H.B. J.A. is funded by the Deanship of Scientific Research at Northern Border University, Arar, KSA through the project number NBU-SAFIR-2024.

## Notes

### Competing Interest Statement

The authors have declared no competing interest.

