## Supplementary Information for "Accessibility of Telomeric Overhangs to Stabilizing Small-Molecule Ligands"

**DNA Sequences:** The sequences of strands that were used to create the partial duplex DNA (pdDNA) constructs and the figures they are employed in are given in Table S1.

| Strands | Sequence in 5′–3′ | Stem | Used in Figures |
|---|---|---|---|
| nG-Tract | TGGCGACGGCAGCGAGGCTTAGGGTTAGGG $(TTAGGG)_n$ TTAG | Stem-28-Cy3@bp4 or Stem-28 | Figs. 2, 3, S2, S3, S4 |
| Stem-28-Cy3@bp4 | CTAA/iCy3/CCCTAAGCCTCGCTGCCGTCGCCA-biotin | | Figs. 2, 3, S2, S3B, S4, S5A, S6A |
| Stem-28 | CTAACCCTAAGCCTCGCTGCCGTCGCCA-biotin | | Fig. 3D, S5B, S6B |
| Quencher | CGA GGC TTA GGG /3BHQ_2/ | | Figs. 2, 3, S2, S3, S4 |
| nG-Tract : 9-nt Spacer | TGGCGACGGCAGCGAGGC $(TTAGGG)_4$ $[(TTA)_3$ $(GGG\ TTA)_4$ $(TTA)_2]_n$ $(GGGTTA)_4$ G | Stem-28-Cy3@bp4 or Stem-28 | Fig. 3 |
| 3L1L-nRep: 3-nt Spacer | TGGCGACGGCAGCGAGGCTTA $[(GGGT)_3$ $GGGTTA]_n$ | Stem-28-Cy3, or Stem-28 | Fig. S5, S6 |
| 3L1L-nRep: 6-nt Spacer | TGGCGACGGCAGCGAGGCTTA $[(GGGT)_3$ GGG $TTATTA]_{m-1}$$[(GGGT)_3$ GGGTTA] | Stem-28-Cy3, or Stem-28 | Fig. S5 |
| 3L1L-nRep: 9-nt Spacer | TGGCGACGGCAGCGAGGCTTA $[(GGGT)_3$ GGG $TTATTATTA]_{m-1}$ $[(GGGT)_3$ GGGTTA] | Stem-28-Cy3, or Stem-28 | Fig. S5 |

**Table S1**: Sequences used for constructing pdDNA constructs. Purple bases on the long strands pair with the purple bases of the stem strands (Stem-28-Cy34bp, Stem-28, Stem-28-Cy3, or Stem-28) to form the duplex region. Nucleotides shown in green denote the single-stranded overhang. Subscripts indicate the number of repeats, e.g., $(TTAGGG)_2$ corresponds to TTAGGGTTAGGG. The purple nucleotides in the Quencher strand are complementary to the Stem-28-Cy3-bp4 segment and position a /3BHQ_2/ ("black hole quencher") group directly across from Cy3 in the duplex, enabling efficient fluorescence quenching. The 'nG-Tract: 9-nt Spacer' constructs contain 9-nt spacers d(TTATTATTA) inserted at defined positions within the telomeric repeat sequence such that they separate consecutive GQs. The '3L1L-nRep' constructs contain $n$ repeats of GGGT sequence. The $m$ GQs in these constructs are separated from each other by 3-nt d(TTA), 6-nt d(TTATTA), or 9-nt d(TTATTATTA) spacers, as specified in the table.

| | Number of Molecules | | | | | | |
|---|---|---|---|---|---|---|---|
| | 1 STEP | 2 STEPS | 3 STEPS | 4 STEPS | 5 STEPS | 6 STEPS | TOTAL |
| 4GTr | 178 | 30 | 0 | 0 | 0 | 0 | 208 |
| 6GTr | 170 | 64 | 0 | 0 | 0 | 0 | 234 |
| 8GTr | 196 | 26 | 10 | 6 | 0 | 0 | 238 |
| 10GTr | 143 | 50 | 15 | 10 | 0 | 0 | 218 |
| 12GTr | 294 | 86 | 36 | 13 | 3 | 2 | 434 |
| 14GTr | 136 | 82 | 56 | 33 | 9 | 3 | 319 |
| 16GTr | 102 | 76 | 46 | 30 | 10 | 6 | 270 |
| 18GTr | 80 | 90 | 56 | 40 | 10 | 8 | 284 |
| 20GTr | 74 | 82 | 54 | 38 | 14 | 8 | 270 |
| 22GTr | 56 | 84 | 54 | 38 | 12 | 8 | 252 |
| 24GTr | 56 | 98 | 76 | 50 | 16 | 10 | 261 |
| 26GTr | 28 | 57 | 43 | 29 | 10 | 6 | 173 |

**Table S2**: Number of molecules presented in histograms in Fig. 2A and Fig. S2.

| | Number of Molecules | | | | | | |
|---|---|---|---|---|---|---|---|
| | 1 STEP | 2 STEPS | 3 STEPS | 4 STEPS | 5 STEPS | 6 STEPS | TOTAL |
| 6GTr: 3-nt | 282 | 60 | 0 | 0 | 0 | 0 | 342 |
| 10GTr: 3-nt | 162 | 54 | 18 | 10 | 0 | 0 | 244 |
| 14GTr: 3-nt | 178 | 64 | 50 | 28 | 8 | 4 | 332 |
| 18GTr: 3-nt | 82 | 94 | 52 | 38 | 10 | 8 | 284 |
| 22GTr: 3-nt | 66 | 72 | 46 | 36 | 12 | 6 | 238 |
| 6GTr: 9-nt | 98 | 60 | 0 | 0 | 0 | 0 | 158 |
| 10GTr: 9-nt | 72 | 44 | 31 | 19 | 0 | 0 | 166 |
| 14GTr: 9-nt | 64 | 48 | 36 | 18 | 10 | 6 | 182 |
| 18GTr: 9-nt | 49 | 54 | 42 | 34 | 23 | 12 | 214 |
| 22GTr: 9-nt | 41 | 58 | 43 | 29 | 20 | 11 | 202 |

**Table S3**: Number of molecules presented in histograms in Fig. 3B

| Overhang length | T-Test |
|---|---|
| 6GTr | $t(4)= 4.81, p=0.009$ |
| 10GTr | $t(4)= 15.74, p<0.001$ |
| 14GTr | $t(3)= 4.00, p=0.028$ |
| 18GTr | $t(2)= 7.75, p=0.016$ |
| 22GTr | $t(3)= 4.49, p=0.021$ |

**Table S4**: T-Test analysis illustrating the statistical significance of the differences in the average number of bound molecules between the 3-nt Spacer constructs and the 9-nt Spacer constructs that contain the same number of G-tracts (data in Fig. 3B).

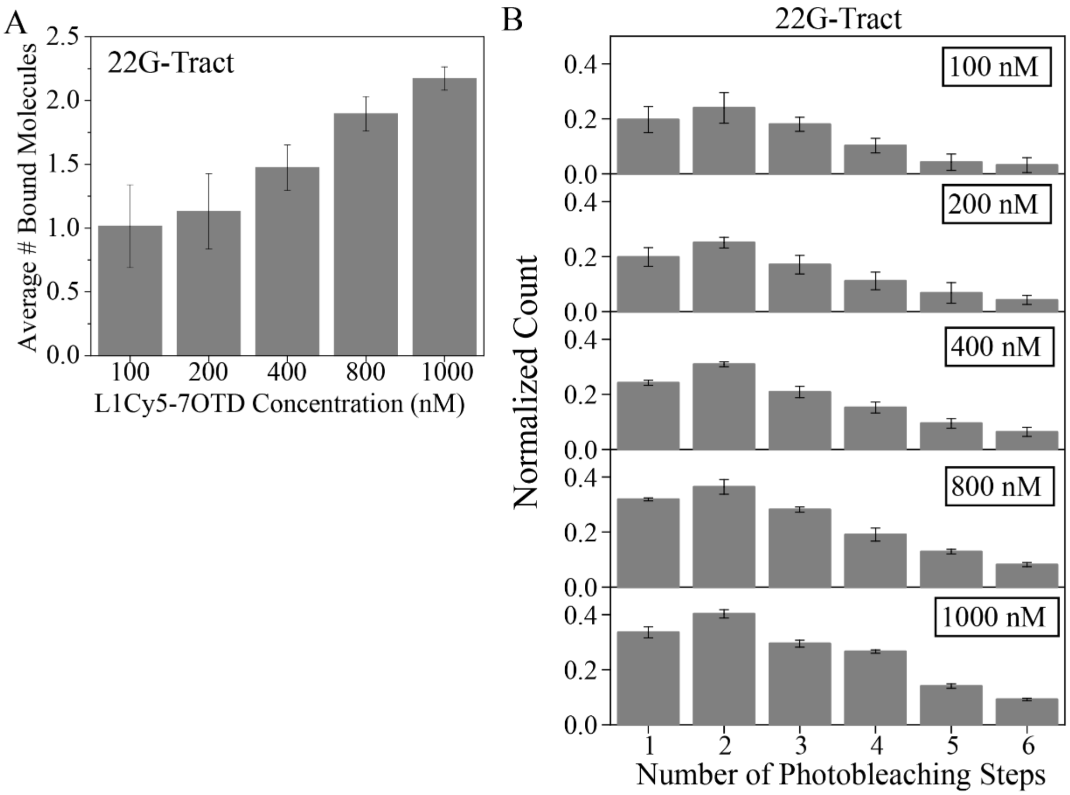


**Figure S1.** Photobleaching step counting assay to investigate impact of L1Cy5-7OTD concentration on binding stoichiometry. (A) Average number of bound molecules for the 22GTr as a function of L1Cy5-7OTD concentration. Increasing L1Cy5-7OTD concentration from 100 nM to 1000 nM resulted in a ~2.2-fold increase in average number of bound ligands (1.01 ± 0.33 to 2.17 ± 0.09). A one-way ANOVA revealed a significant effect of concentration on the average number of bound molecules, $F(4, 16) = 1.48, p < .001$. (B) Two-dimensional histograms showing the normalized distributions of photobleaching steps for each concentration, based on single-molecule traces. Concentrations beyond 1000 nM could not be reliably analyzed due to large background. Error bars represent the standard error from independent measurements.

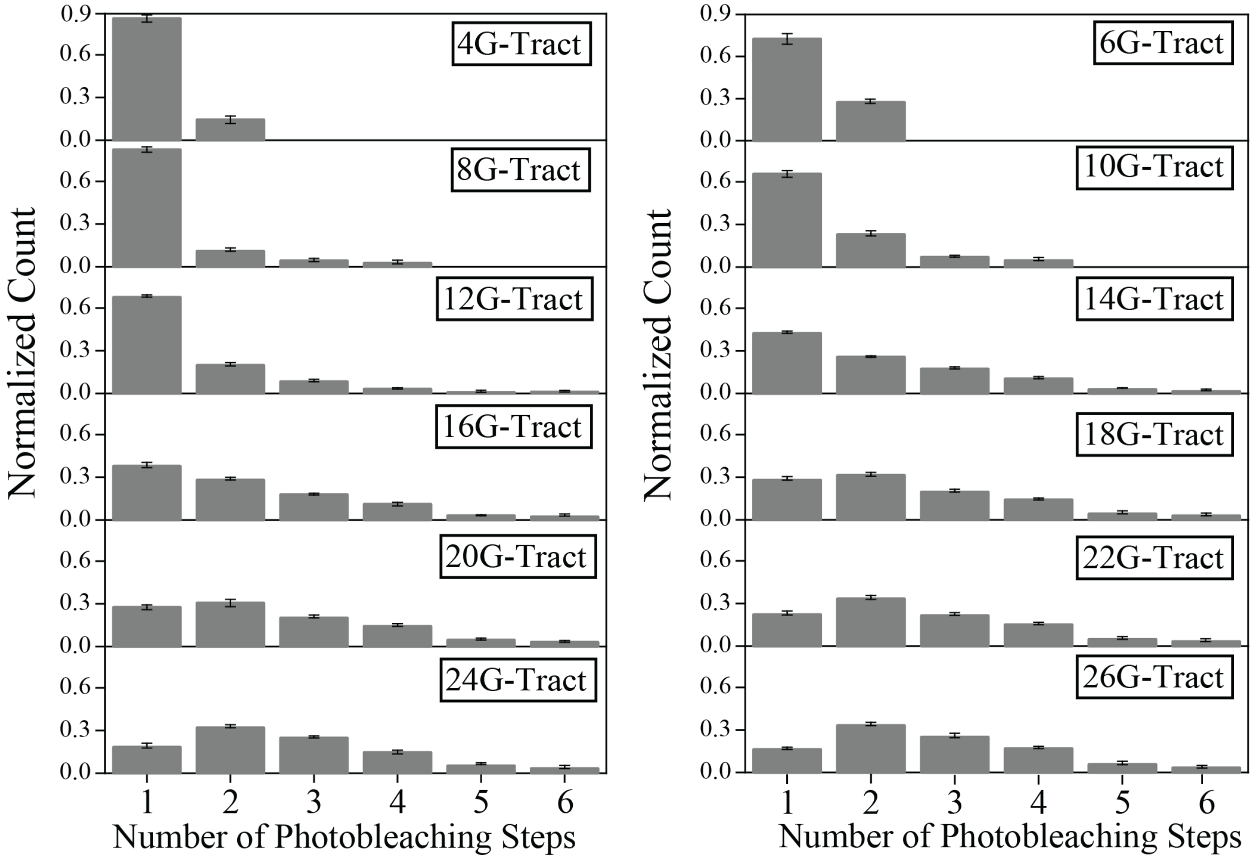


**Figure S2**. Photobleaching step distributions of L1Cy5-7OTD binding to telomeric constructs with varying overhang lengths. Two-dimensional histograms show the normalized distributions of binding events based on the number of discrete photobleaching steps observed in single-molecule traces. The 3D versions of these data are presented in Fig. 2. Left panel shows constructs containing exact numbers of G-tracts ($[4n]$GTr) corresponding to 1-6 G-quadruplexes (4GTr-24GTr), while the right panel shows constructs extended by two additional G-tracts (6GTr-26GTr) ($[4n+2]$GTr). These data do not include the molecules that did not show any L1Cy5-7OTD molecule binding (e.g., zero steps).

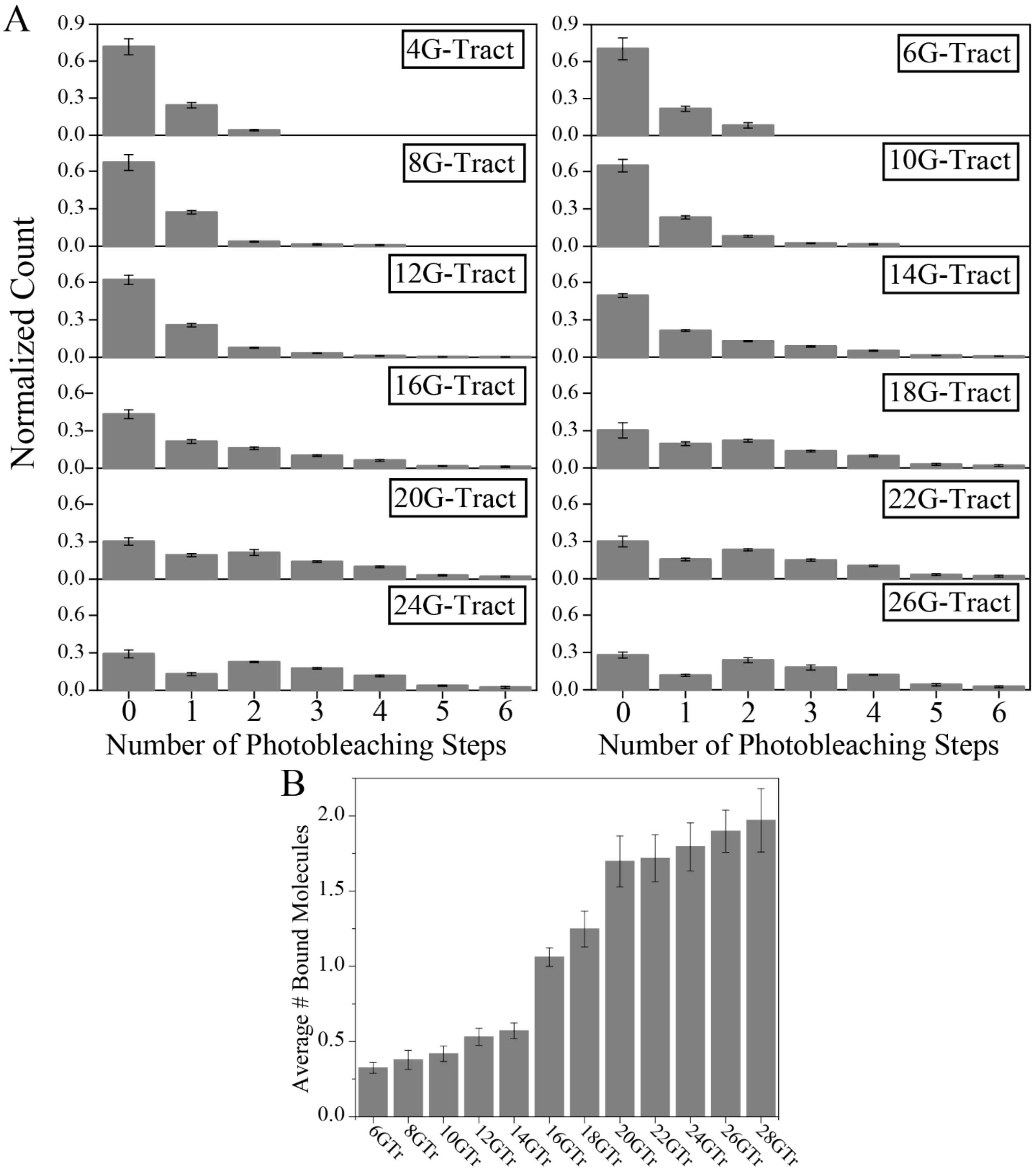


**Figure S3.** Photobleaching step analysis that includes constructs showing zero photobleaching steps. (A) Photobleaching step distributions of L1Cy5-7OTD binding to telomeric constructs when DNA constructs that did not show any L1Cy5-7OTD binding (zero-step events) are included in the analysis. (B) Average number of bound L1Cy5-7OTD molecules per DNA construct when zero step events are included. Bars represent mean values from different measurements (n=5-10); error bars indicate the standard error associated with these measurements.

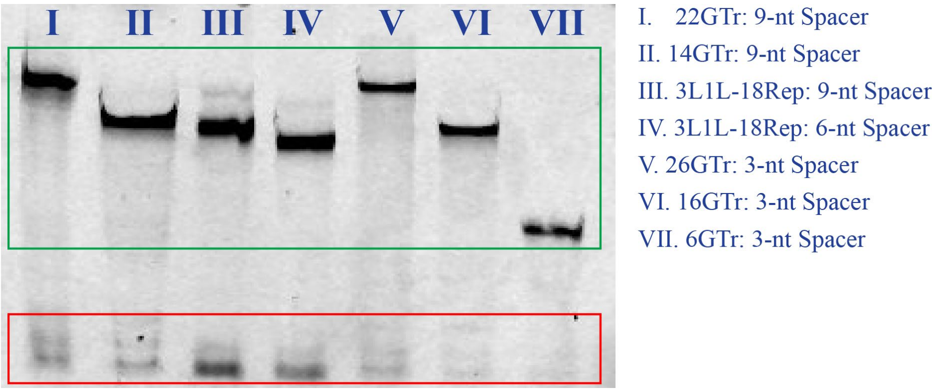


**Figure S4**. Native polyacrylamide gel electrophoresis (PAGE) examination of pdDNA hybridization efficiency. Telomeric overhang constructs containing different numbers of G-tracts and spacers length were annealed with a Cy3-labeled 28-nt complementary stem strand and resolved by native PAGE. The upper bands (green box) represent hybridized pdDNA, while the lower bands (red box) correspond to unhybridized stem strands (just single-stranded DNA with a Cy3). Quantification of band intensities was used to estimate the fraction of unhybridized DNA for each construct: 22GTr: 9-nt Spacer (15%), 14GTr: 9-nt Spacer in 1 M KCl (29%), 3L1L-18Rep: 9-nt Spacer (36%), 3L1L-18Rep: 6-nt Spacer (35%), 26GTr: 3-nt Spacer (28%), 16GTr: 3-nt Spacer (25%), and 6GTr: 3-nt Spacer (5.7%). Across all constructs, the majority of DNA constructs migrated in the duplex band, demonstrating efficient hybridization, while the amount of unhybridized ssDNA fraction varied with sequence composition and spacer length. The 14GTr: 9-nt Spacer construct was annealed in 1 M KCl (unlike all other constructs which were annealed in 150 mM KCl) to test the impact of ion concentration but the hybridization efficiency did not increase with this change in ion concentration.

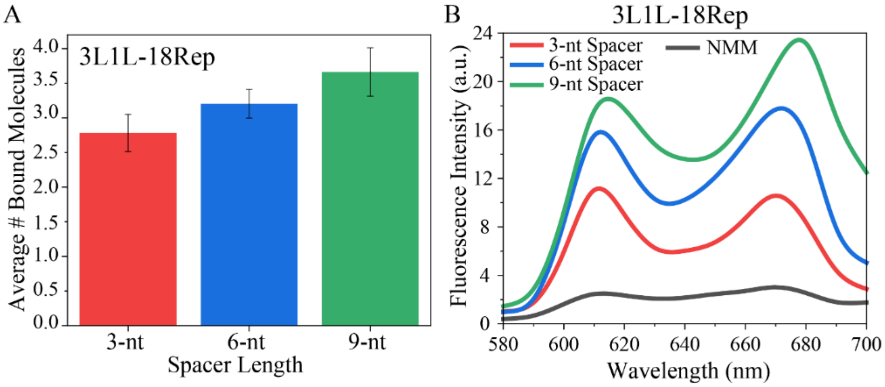


**Figure S5**. L1Cy5-7OTD (single molecule fluorescence) and NMM (bulk fluorescence) binding measurements on 3L1L constructs with 18 repeats of d(GGGT) but varying spacer length between consecutive GQs, which are separated from each other by 3-nt d(TTA), 6-nt d(TTATTA), or 9-nt d(TTATTATTA) spacers. (A) Average number of bound L1Cy5-7OTD molecules ($\langle n_b \rangle$) for these constructs was determined using single molecule fluorescence measurements. Mean occupancies (± standard error) increased from 2.78 ± 0.27 (3-nt-spacer, red) to 3.21 ± 0.21 (6-nt-spacer, blue) and 3.66 ± 0.35 (9-nt-spacer, green) indicating that longer spacers support progressively higher binding stoichiometry. A one-way ANOVA revealed a significant effect of spacer length on the average number of bound molecules, $F(2, 11) = 8.16, p < .001$. (B) Fluorescence spectra of N-methyl mesoporphyrin IX (NMM) before (black) and after binding to 3L1L-18Rep constructs with varying spacer length. Colored traces correspond to different spacer lengths: red, 3-nt spacer; blue, 6-nt spacer; green, 9-nt spacer. The NMM emission peak at ~610 nm increased from 2.4 a.u. for NMM alone to 11.0, 15.5, and 17.5 a.u. for the 3-nt, 6-nt, and 9-nt spacer constructs, respectively, corresponding to ~4.6-fold, ~6.5-fold, and ~7.3-fold enhancement, respectively.

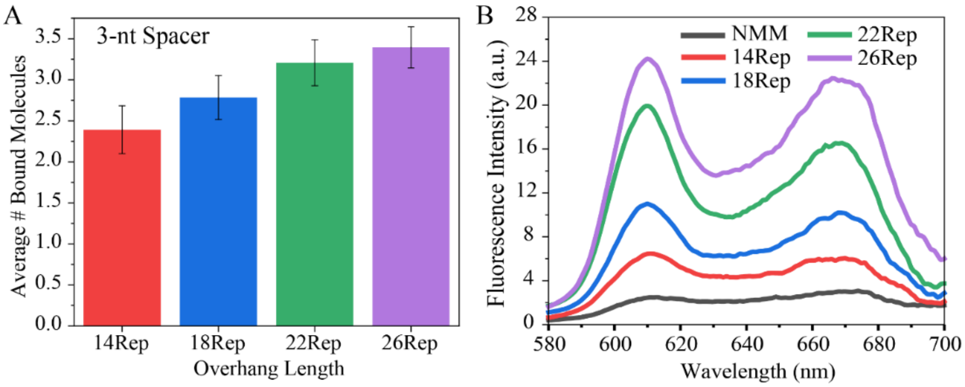


**Figure S6**. L1Cy5-7OTD (single molecule fluorescence) and NMM (bulk fluorescence) binding analysis for 3L1L constructs with varying overhang length. (A) Average number of bound L1Cy5-7OTD molecules ($\langle n_b \rangle$) for 3L1L constructs with 3-nt-spacer and varying overhang length. Mean occupancies (± standard error) were 2.39 ± 0.29 (14GTr), 2.78 ± 0.27 (18GTr), 3.21 ± 0.68 (22GTr), and 3.39 ± 0.25 (26GTr), demonstrating a systematic increase in binding stoichiometry with overhang length. A one-way ANOVA revealed a significant effect of overhang length on the average number of bound molecules, $F(3, 15) = 17.43, p < .001$. (B) Fluorescence spectra of N-methyl mesoporphyrin IX (NMM) before and after binding to 3L1L constructs of varying overhang length (all with 3-nt spacers between GQs). Colored curves correspond to individual constructs: black: NMM-only; red: 14Rep; blue: 18Rep; green: 22Rep; and purple: 26Rep. NMM emission spectra showed markedly higher fluorescence intensity in the presence of 3L1L GQs compared to NMM alone. The NMM fluorescence emission signal based on the peak amplitude at 610 nm increased from 2.4 a.u. for NMM alone to 6.4, 11.0, 19.9, and 24.2 a.u. for the 14Rep, 18Rep, 22Rep, and 26Rep constructs, respectively, corresponding to ~2.7-fold, ~4.6-fold, ~8.3-fold, and ~10.1-fold enhancements, respectively. These results indicate that longer overhangs accommodate binding of a larger number of NMM.

**Lattice Model for Cooperative GQ Folding and Ligand Binding**

**Transfer Matrix:** The transfer matrix, $T_i$ describes the statistical weight of the $i^{th}$ repeat given that $(i+1)^{th}$ repeat is in state $\gamma$. In the absence of ligands, a repeat can either be unfolded (1) or be one of four consecutive repeats in a GQ (labeled $\ell^{(1-4)}$, where $1-4$ refer to the position of the repeat within the folded GQ starting from the $5'$-end). Small molecules can bind to the top interface (labeled $t^{(1-4)}$), to the bottom interface (labeled $b^{(1-4)}$), or to both the top and bottom interfaces (labeled $(bt)^{(1-4)}$) of a folded GQ.

The transfer matrix is therefore a $17 \times 17$ matrix with indices labeled by the states as:

$$1\ \ell^{(1)}\ \ell^{(2)}\ \ell^{(3)}\ \ell^{(4)}\ b^{(1)}\ b^{(2)}\ b^{(3)}\ b^{(4)}\ t^{(1)}\ t^{(2)}\ t^{(3)}\ t^{(4)}\ (bt)^{(1)}\ (bt)^{(2)}\ (bt)^{(3)}\ (bt)^{(4)}$$

Non-zero elements of the transfer matrix $T_i(m,n)$ and the special characteristic of the junction (between double and single stranded telomeres) and the free $3'$-end are described below.

- **Unfolded repeat ($\boldsymbol{m=1}$):** Unfolded repeats have a weight of 1, and can be followed by an unfolded repeat, or the first repeat that is a folded GQ (with or without ligands):

$$T_i(1,\gamma) = 1, \text{ where } \gamma = 1, \ell^{(1)},\ b^{(1)},\ t^{(1)},\ (bt)^{(1)}$$

- **Folded GQ, unbound by a ligand ($\boldsymbol{m = \ell^{(1-4)}}$)**: Folded GQs are four consecutive repeats with a statistical weight of $K_\mathrm{f}$. We associate this weight with the first repeat in the GQ,

$$T_i(\ell^{(1)},\ell^{(2)}) = K_\mathrm{f} \text{ and } T_i(\ell^{(2)},\ell^{(3)}) = T_i(\ell^{(3)},\ell^{(4)}) = 1.$$

The last repeat in the GQ needs to be treated differently as it can be followed by an unfolded repeat or by the first repeat in a neighboring GQ. The neighboring GQs have an additional cooperativity statistical weight ($K_c$), which we associate the cooperativity with the last repeat in the GQ. We maintain the same $K_c$ regardless of whether the neighboring GQ is without a ligand, with a top-bound, bottom-bound, or top-and-bottom bound ligands. These are expressed as:

$$T_i(\ell^{(4)},1) = 1 \text{ and } T_i(\ell^{(4)},\gamma) = K_\mathrm{c}, \text{ where } \gamma = \ell^{(1)},\ b^{(1)},\ t^{(1)},\ (bt)^{(1)}.$$

- **Folded GQ, ligand bound to bottom interface ($\boldsymbol{m = b^{(1-4)}}$):** A statistical weight of $b$ is associated with a small molecule bound to the bottom interface of a folded GQ giving an overall weigth of $K_\mathrm{f} \times b$. The matrix elements are otherwise similar to unbound folded GQ:

$$T_i(b^{(1)},\ b^{(2)}) = K_\mathrm{f} \times b, \text{ and } T_i(b^{(2)},\ b^{(3)}) = T_i(b^{(3)},\ b^{(4)}) = 1.$$

The last repeat in the GQ can be followed by an unfolded repeat or by the first repeat in a neighboring GQ that is unbound, or bound by a top, bottom, or top-and-bottom ligands. Neighboring GQs have an additional cooperativity statistical weight $K_\mathrm{c}$ which we associate the cooperativity with the last repeat in the GQ. These are expressed as:

$$T_i(b^{(4)},1) = 1 \text{ and } T_i(b^{(4)},\gamma) = K_\mathrm{c}, \text{ where } \gamma = \ell^{(1)},\ b^{(1)},\ t^{(1)},\ (bt)^{(1)}.$$

- **Folded GQ, molecule bound to top interface ($\boldsymbol{m} = \boldsymbol{t}^{(\mathbf{1-4})}$):** A statistical weight of $t$ is associated with a small molecule bound to the top interface of a folded GQ giving an overall weigth of $K_\text{f} \times t$,

$$T_i(\, t^{(1)},\ t^{(2)}) = K_\text{f} \times t, \text{ and } T_i(\, t^{(2)},\ t^{(3)}) = T_i(\, t^{(3)},\ t^{(4)}) = 1.$$

The last repeat in the GQ with molecule bound to the top surface can be followed by an unfolded repeat or by the first repeat in a neighboring GQ that is unbound, or bound by a top, bottom, or top-and-bottom ligands. Neighboring GQs have an additional cooperativity statistical weight $K_\text{c}$, unless followed by a folded GQ with a molecule bound to the bottom interface. These are expressed as:

$$T_i(\, t^{(4)}, 1) = 1 \text{ and } T_i(\, t^{(4)},\ \ell^{(1)}) = T_i(\, t^{(4)},\ t^{(1)}) = K_c.$$

An additional statistical weight of $\omega_\text{tb}$ is associated with a top bound GQ followed by a GQ with a molecule bound to the bottom interface:

$$T_i(\, t^{(4)},\ b^{(1)}) = T_i(\, t^{(4)},\ (bt)^{(1)}) = K_c\omega_\text{tb}.$$

- **Folded GQ, molecule bound to top-and-bottom interface ($\boldsymbol{m} = (\boldsymbol{bt})^{(\mathbf{1-4})}$):** A statistical weight of $(bt)$ is associated with small molecules bound to the top-and-bottom interfaces of a folded GQ giving an overall weight of $K_\text{f} \times t \times b$, which is associated with the first repeat:

$$T_i(\, (bt)^{(1)},\ (bt)^{(2)}) = K_\text{f} \times t \times b, \text{ and } T_i(\, (bt)^{(2)},\ (bt)^{(3)}) = T_i(\, (bt)^{(3)},\ (bt)^{(4)}) = 1.$$

The last repeat in this GQ can be followed by an unfolded repeat or by the first repeat in a neighboring GQ that is unbound, or bound by a top, bottom, or top-and-bottom ligands. Neighboring GQs have an additional cooperativity statistical weight $K_\text{c}$ (associated with this last repeat), unless followed by a folded GQ with a molecule bound to the bottom interface:

$$T_i(\, (bt)^{(4)}, 1) = 1 \text{ and } T_i(\, (bt)^{(4)},\ \ell^{(1)}) = T_i(\, (bt)^{(4)},\ t^{(1)}) = K_\text{c}.$$

An additional statistical weight of $\omega_\text{tb}$ is associated with a top bound GQ followed by a GQ with a molecule bound to the bottom interface, which results in:

$$T_i(\, (bt)^{(4)},\ b^{(1)}) = T_i(\, (bt)^{(4)},\ (bt)^{(1)}) = K_c\omega_\text{tb}.$$

- **Junction Region:** G-quadruplexes are destabilized at the junction region and have different small molecule binding affinities. Therefore, the first repeat at the interface of the junction region is described with a different transfer matrix $T_1$ in which $K_f$, $t$, and $b$ are replaced by $K_{f1}$, $t_1$, and $b_1$, respectively. Similarly, a GQ starting at the second repeat is described by a transfer matrix $T_2$ in which $K_f$ is replaced by $K_{f2}$.

- **The 3′-end:** At the 3′-end, GQs ending at the next-to-last or the last repeat also have different stabilities compared to a GQ in the interior of the telomere sequence. To account for this, the $K_f$ in the transfer matrix $T_{N-4}$ is replaced $K_{fN-1}$. Similarly, the $K_f$ in the transfer matrix $T_{N-3}$ is replaced $K_{fN}$. The transfer matrices for $T_{N-2}, T_{N-1}, and\ T_N$ are unchanged since the full weight of the GQ is associated with its first repeat of the GQ, which corresponds to $T_{N-4}$ for a GQ ending in next-to-last repeat and $T_{N-3}$ for a GQ ending in the last repeat.

**The Partition Function:** The partition function can be written as:

$$Z = V_1^T T_1 T_2 \cdots\cdots T_{N-1} T_N V_N$$

This expression is equivalent to summing over all possible configurations of the system while row vector $V_1^T$ and column vector $V_N$ enforce boundary conditions of the available states for the first and last repeats, respectively. The non-zero elements of $V_1$ are $V_1(\gamma) = 1$, for $\gamma = 1,\ \ell^{(1)},\ b^{(1)},\ t^{(1)},\ (bt)^{(1)}$, which are the possible states for the first repeat. Similarly, the statistical weight for the last repeat can be written as $\tilde{V}_N(\gamma) = 1$ for $\gamma = 1, \ell^{(4)}, b^{(4)}, t^{(4)}, (bt)^{(4)}$, i.e., the last repeat can either be unfolded or the fourth repeat of a folded GQ that can be in one of familiar four states (not bound by a ligand, bound in the top face, bottom face or top-and-bottom faces). This is the first column of $T_N$, so we can write this vector of weights as $\tilde{V}_N = T_N V_N$, where $V_N$ is a vector whose only non-zero element is $V_N(1) = 1$.

Many thermodynamic quantities can be directly computed with the help of transfer matrices. For example, the probability that the $i^{th}$ repeat has particular property $\beta$ (such as being unfolded) is:

$$p_\beta(i) = \frac{1}{Z} V_1^T T_1 T_2 \cdots \tilde{T}(\beta) T_{i+1} \cdots T_{N-1} T_N V_N,$$

where $\tilde{T}(\beta)$ is a matrix consistent with the property $\beta$. For example, to find the probability that a repeat is unfolded, then $T_i$ is replaced with a matrix $\tilde{T}$ which has non-zero elements only in the row corresponding to the unfolded state.

**Determining binding coefficients:** Calculating the probability of having $n_\mathrm{b}$ bound ligands for a construct of $N$ repeats, $P_N(n_\mathrm{b})$ is not particularly convenient with transfer matrices directly. To calculate $P_N(n_\mathrm{b})$, we generated the weights of the states with transfer matrices using symbolic parameters $t$, $b$, and $(bt)$ and collected the relevant terms consistent with $n_\mathrm{b}$ bound ligands using Mathematica.

The values for the statistical weights of the GQ folding reported in the paper are chosen as a representative set consistent with the localization pattern measured in our earlier publication, Shiekh et al. (PNAS, 2022). To find the values for the binding parameters $t$, $b$, and $\omega_{tb}$, we performed a least squared fit to the experimentally measured probability of the number of bound ligands, $P_N^{\mathrm{meas}}(n_\mathrm{b})$, while keeping other parameters fixed.

Let $Q$ be the set of repeat lengths we fit where $Q = \{6, 8, 10, 12, 14, 16, 18, 20, 22, 24, 26\}$. For each $N \in Q$, let $B(N)$ be the set of allowed bound molecule counts: $B(N) = \{0,1, \dots, n_b^{\max}\}$. That is, we minimized

$$A = \sum_{\substack{n_b \in B(N) \\ N \in Q}} (P_N^{meas}(n_b) - P_N^{model}(n_b, t, b, \omega_{tb}))^2$$

where $P_N^{\text{model}}(n_b, t, b, \omega_{tb})$ is the calculated probability of number of bound ligands with all other parameters fixed. To optimize A with respect to $t$, $b$, and $\omega_{tb}$, we compute symbolic partial derivatives $\partial A/\partial t$, $\partial A/\partial b$, and $\partial A/\partial \omega_{tb}$ in Mathematica and then minimize $A$ using steepest descent. Further, we report the root-mean-squared error (RMSE), $\sqrt{A/M}$ , where $M$ is the total number of $\{N, n_b\}$ pairs used in the summation for $A$.

It turns out that $\omega_{tb}$ only influences the binding distribution for larger N because there are more opportunities for top-bottom adjacent binding pairs. Therefore, we split the optimization into two parts: optimization of $A$ for $t$ and $b$ with $\omega_{tb}$ fixed for the whole data set, and optimization of $A$ with respect to $\omega_{tb}$ with $t$ and $b$ fixed for the data set restricted to $N = 22, 24, 26$. These two optimizations were solved self-consistently.

The model is symmetric with respect to top vs. bottom binding, so unconstrained minimization of $A$ with $\omega_{tb}$ fixed yields the optimum at approximately equal $t$ and $b$, specifically $t = b = 0.4$. Since the binding to the bottom interface is known to be weaker than to the top, we investigate the binding coefficients constrained to $t = \alpha b$ with $\alpha > 0$ and optimize over $b$. Along the constrained line, the directional derivative is

$$\frac{dA}{db} = \frac{\partial A}{\partial b} + \alpha \frac{\partial A}{\partial t}.$$

We use this to update $b$ by steepest descent to find the optimum $b$. Then, with $b$ held fixed, we optimize $\omega_{tb}$ using steepest descent with $\partial A/\partial \omega_{tb}$. We iterate these two optimizations until $b$ *and* $\omega_{tb}$ converge to $b_{\text{opt}}(\alpha)$ and $\omega_{tb,\text{opt}}(\alpha)$.

Under this constraint, the method gives roughly $t + b = 0.8$, or $b_{\text{opt}}(\alpha) = \frac{0.8}{1+\alpha}$. As shown in Fig. S7, the binding cooperativity between successive top and bottom binding molecules is always negative and quite destabilizing. Interestingly, there is a shallow minimum at $\alpha = 4$. Increasing asymmetry between $t$ and $b$ demands progressively weaker negative cooperativity ($\omega_{tb}$ increasing towards 1) to reproduce the measured occupancies.

The RMSE as a function of $\alpha$ is shown in Fig. S8. The fits demonstrate good quality, regardless of the value of $\alpha$. We select $\alpha = 3$ as the $t/b$ ratio to compare with the experiment as an illustration. With this choice, the fitted model gives RMSE = $\pm 0.055$ with $b_{opt} = 0.20$, $t_{opt} = 0.60$, and $\omega_{tb} = 0.03$ .

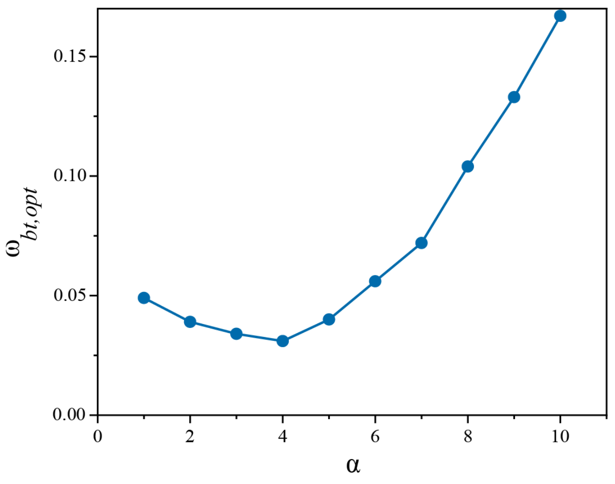


**Figure S7.** Optimal molecular binding cooperativity vs. asymmetry ratio alpha. Best-fit $\omega_{tb,\mathrm{opt}}$ as a function of the imposed asymmetry $\alpha$ in the line-search constraint $t = \alpha b$. The curve shows a shallow minimum near $\alpha \approx 4$ followed by a monotonic rise, reaching $\omega_{tb,\mathrm{opt}} \approx 0.16$ at $\alpha = 10$.

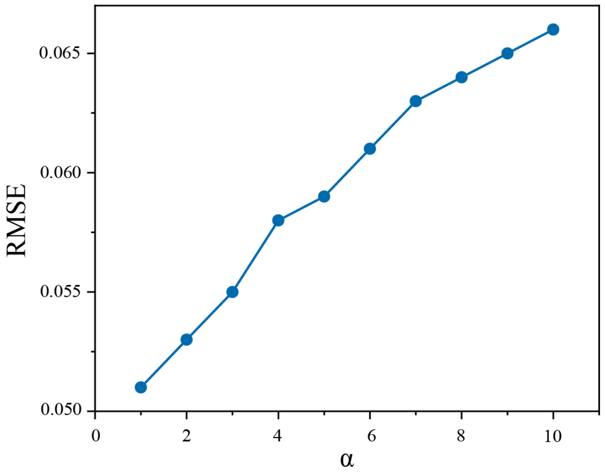


**Figure S8.** Fit quality vs. asymmetry ratio $\alpha$. Root-mean-squared error, RMSE = $\sqrt{A/M}$, is plotted as a function of the constraint ratio $\alpha$. RMSE is computed over all $(N, n_b)$ pairs included in the fit ($Q = \{6,8, \ldots ,26\}$; $M$ = total number of data pairs).

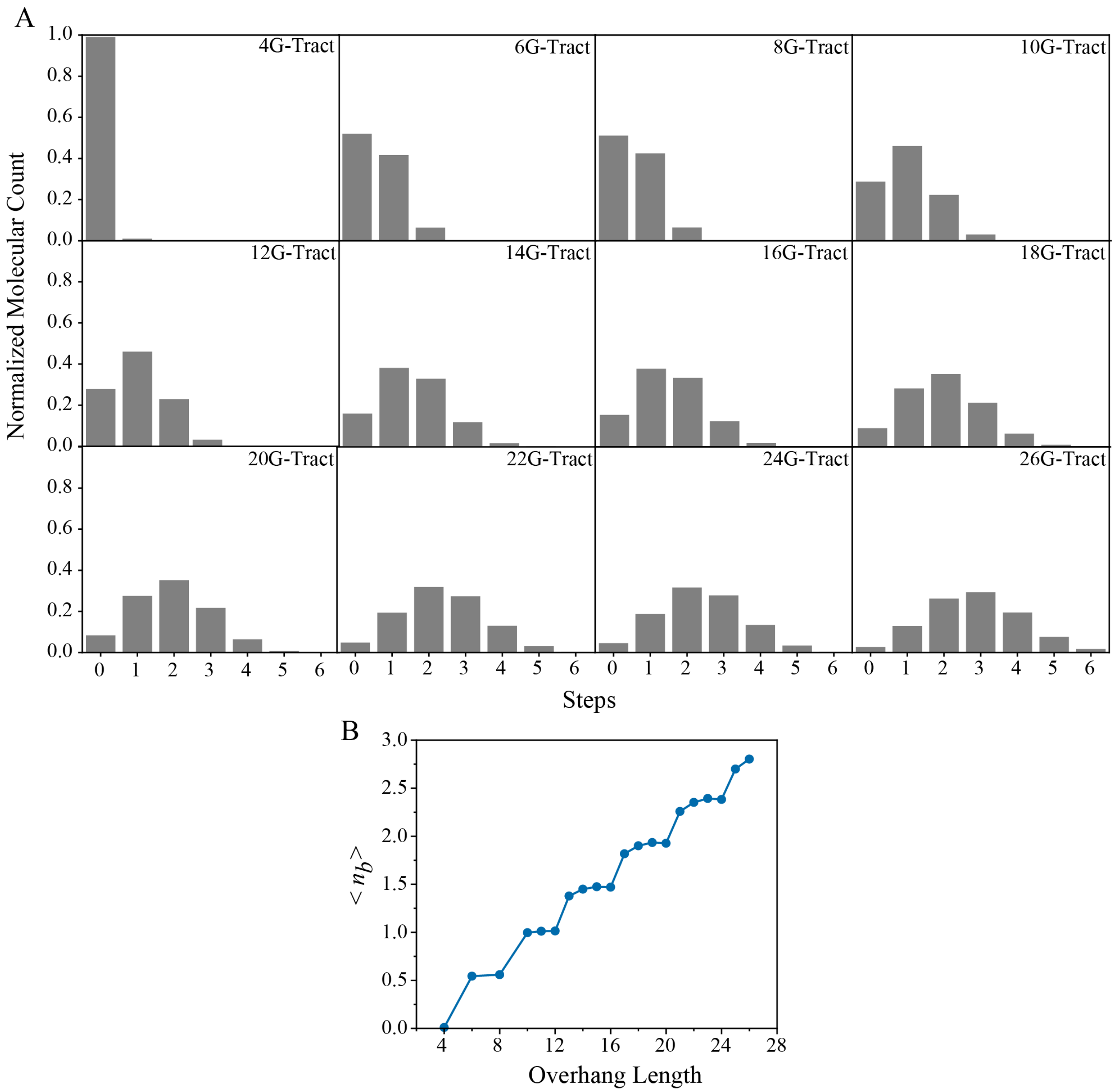


**Figure S9.** Normalized molecular count distributions **(A),** and average molecular occupancy **(B)** as a function of telomeric overhang length including constructs with no small molecule binding (zero steps included). Short overhangs favor 1-2 bound molecules, whereas longer overhangs accommodate up to 6. This is similar to Fig. 4B with the average number of bound molecules calculated without excluding the unbound population, $P_N(0)$.

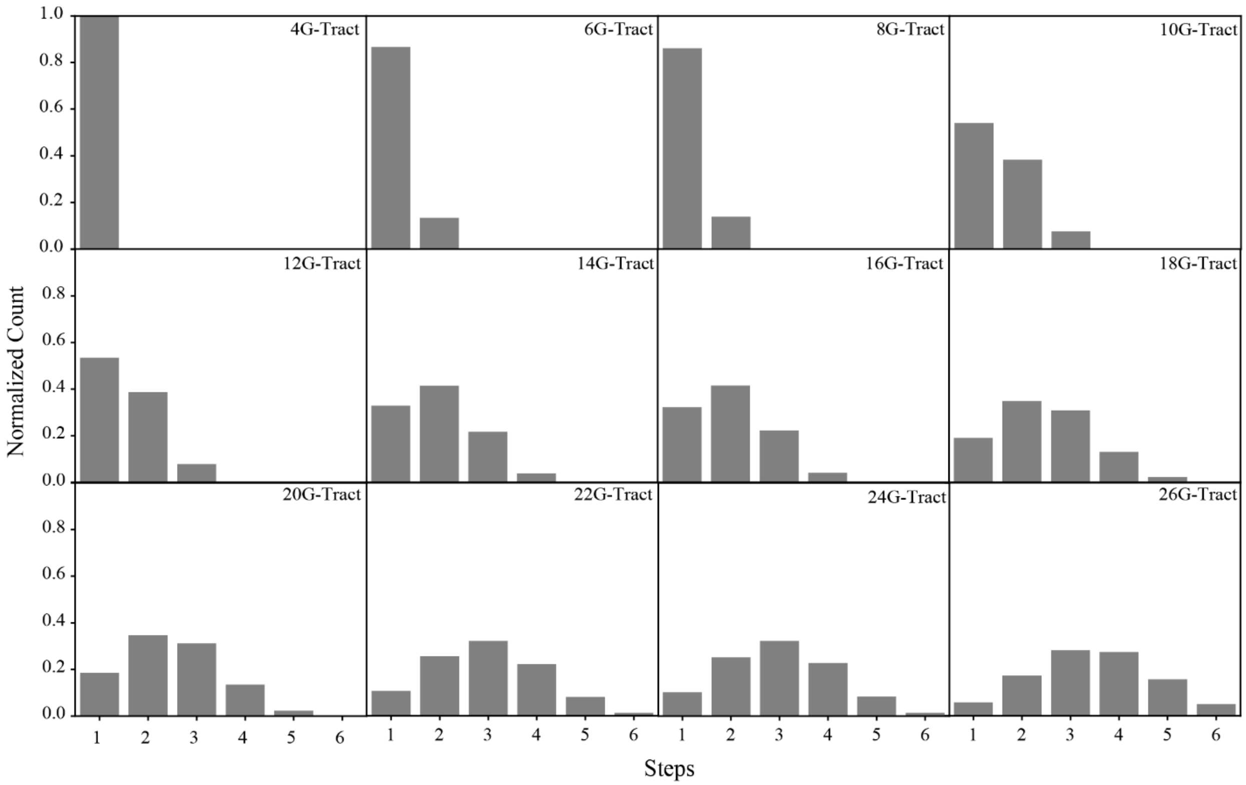


**Figure S10.** Effect of stronger binding constants $b$ and $t$ on occupancy patterns. Increasing the binding constants $b$ and $t$ ($b = 0.3$ , $t = 0.9$) causes the model to place the maximum of the binding distribution at multi-ligand states ($n_b > 1$) for shorter overhangs, whereas the experiments show such maxima only for longer constructs (18G-Tracts and above).

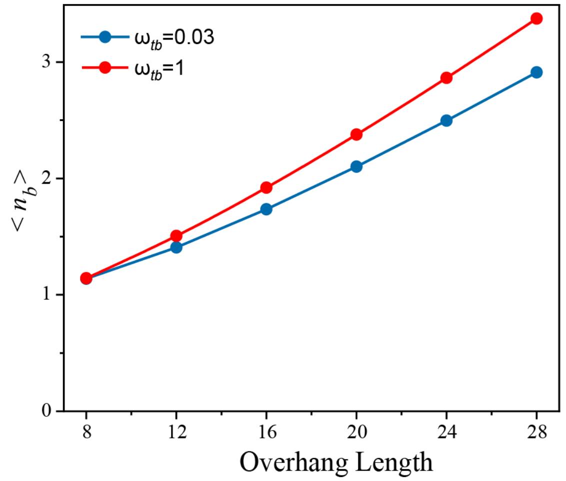


**Figure S11.** Effect of inter-ligand cooperativity $\omega_{tb}$ on mean molecular occupancy. Model-predicted average number of bound molecules $\langle n_b \rangle$ versus overhang length is plotted for two values of adjacent top–bottom cooperativity $\omega_{tb}$. The zero binding events ($P_N(0)$) are excluded in these calculations. Blue line represents negative cooperativity similar to the optimal value we identified in the least square fit while the red line represents no cooperativity. Negative cooperativity suppresses occupancy at all N, and the gap widens with length as adjacent binding opportunities increase. For these calculations, all other folding and binding parameters are held fixed to the values reported in the manuscript.

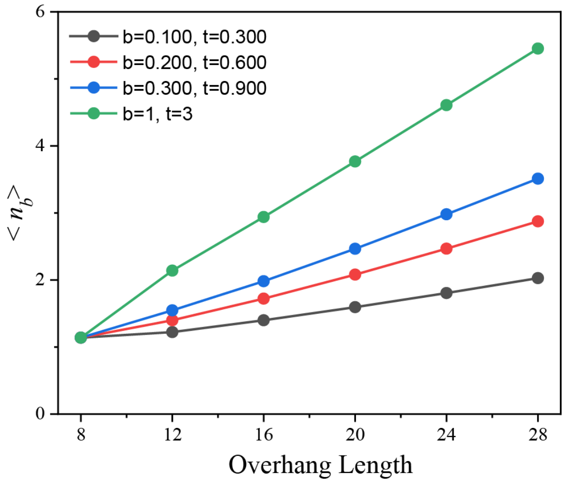


**Figure S12**. Sensitivity of average occupancy to face-specific binding strengths. Average bound-molecule count $\langle n_b \rangle$ versus overhang length (N) are plotted for three (b, t) pairs (bottom-face, top-face weights): (0.1, 0.3), (0.2, 0.6), (0.3, 0.9) and (1.0, 3.0). The zero binding events ($P_N(0)$) are excluded in these calculations. Stronger binding affinities increase both the slope and the value of $\langle n_b \rangle$ with N. Inter-ligand cooperativity $\omega_{tb}$ is negative ($\omega_{tb}$ = 0.03) and all other folding and binding parameters are as those reported in the manuscript.
